# Cross-Frequency Coupling as a Neural Substrate for Prediction Error Evaluation: A Laminar Neural Mass Modeling Approach

**DOI:** 10.1101/2025.03.19.644090

**Authors:** Giulio Ruffini, Edmundo Lopez-Sola, Raul Palma, Roser Sanchez-Todo, Jakub Vohryzek, Francesca Castaldo, Karl Friston

**Affiliations:** Neuroelectrics, Barcelona, Spain; Universitat Pompeu Fabra, Barcelona, Spain; Wellcome Centre for Human Neuroimaging, University College London, London, UK

## Abstract

Predictive coding frameworks suggest that neural computations rely on hierarchical error minimization, where sensory signals are evaluated against internal model predictions. However, the neural implementation of this inference process remains unclear. We propose that cross-frequency coupling (CFC) furnishes a fundamental mechanism for this form of inference. We first demonstrate that our previously described Laminar Neural Mass Model (LaNMM) supports two key forms of CFC: (i) Signal-Envelope Coupling (SEC), where low-frequency rhythms modulate the amplitude envelope of higher-frequency oscillations and (ii) Envelope-Envelope Coupling (EEC), where the envelopes of slower oscillations modulate the envelopes of higher-frequency rhythms. Then, we propose that, by encoding information in signals and their envelopes, these processes instantiate a hierarchical “Comparator” mechanism at the columnar level. Specifically, SEC generates fast prediction-error signals by subtracting top-down predictions from bottom-up oscillatory envelopes, while EEC operates at slower timescales to instantiate gating—a critical computational mechanism for precision-weighting and selective information routing. To establish the face validity and clinical implications of this proposal, we model perturbations of these CFC mechanisms to investigate their roles in pathophysiological and altered neuronal function. We illustrate how, in disorders such as Alzheimer’s disease, disruptions in gamma oscillations following dysfunction in fast-spiking inhibitory interneurons impact Comparator function with an aberrant amplification of prediction errors in the early stages and a drastic attenuation in late phases of the disease. In contrast, by increasing excitatory gain, serotonergic psychedelics diminish the modulatory effect of predictions, resulting in a failure to attenuate prediction error signals (c.f., a failure of sensory attenuation). Collectively, these findings implicate cross-frequency coupling across multiple temporal scales as a key computational mechanism supporting predictive coding and suggest that disruptions in these processes play a central role in disease.

**Highlights:**

- Using an encoding scheme where information is encoded in signals, their envelopes, and envelopes of envelopes, we show how to implement prediction error and precision modulation in a neural mass model through cross-frequency coupling (CFC).
- We use the laminar neural mass model (LaNMM), which integrates Jansen-Rit and pyramidal interneuron gamma (PING) submodels to display fast and slow rhythms and provides mechanisms for a) *Signal-Envelope Coupling (SEC)*, where slow-wave activity modulates the amplitude envelope of fast oscillations (analogous to phase-amplitude coupling), and b) *Envelope-Envelope Coupling (EEC)*, where the envelopes of slower oscillations modulate the envelopes of higher-frequency rhythms.
- We show how to use the LaNMM to implement information-based prediction-error evaluation (as used in Active Inference and Kolmogorov Theory), computing the approximate precision-weighted difference between incoming sensory data (envelopes) and internal model predictions (signals or envelopes).
- We show that using these mechanisms, the Comparator mechanism can operate at multiple levels and timescales, generating fast prediction-error signals (via SEC) and slower gating signals that encode context (e.g., precision) (via EEC).
- Our model provides insights into the physiological and cognitive consequences of mesoscale circuital alterations in the context of predictive coding. First, we study disorders of fast-spiking interneurons, such as Alzheimer’s Disease (AD). In the early stages of AD, error evaluation and precision are disrupted (inflated error and reduced gating/weight of predictions), leading to higher prediction errors. In later stages, prediction errors are suppressed regardless of predictions or their precision.
- Then, we show how serotonergic psychedelics increase the effective weight of inputs and diminish that of predictions, resulting in higher prediction error signals.
- These observations link oscillatory mechanisms and predictive coding alterations, and potentially with the subjective phenomena in each condition—including cognitive decline in AD and hallucinatory states under psychedelics.

## 1 Introduction

### 1.1 Background and Motivation

Predictive coding has become a central framework for understanding cognitive processing. It holds that the brain continuously generates predictions about incoming sensory data and updates these predictions in response to mismatches or prediction errors.^1, 2^

Prediction errors are thought to be passed forward through cortical hierarchies to update higher-level beliefs, while predictions are sent via feedback connections to explain away expected input.^3, 4^ Notably, this bidirectional information flow is reflected in the nature of the extracellular signal—its fundamental features remain consistent across spatial and temporal scales, from microelectrode recordings to large-scale scalp measurements.^5^ This scale-invariance implies that the same principles underwrite local and global dynamics, a property that may facilitate the orchestration of predictive processes under deep (i.e., hierarchical) world models. Neural oscillations are hypothesized to play a key role in this hierarchical inference: higher-frequency activity (particularly in the gamma band, ∼30–100 Hz) is often associated with feedforward signaling of surprise or prediction error, whereas lower-frequency rhythms (alpha/beta bands, ∼8–30 Hz) are linked to top-down predictive signals.^3, 4^ Cross-frequency coupling (CFC), such as the phase or amplitude of slower waves modulating the amplitude of faster oscillations, is thought to coordinate communication between levels of the hierarchy.^6^ This contextual coordination is sometimes cast in terms of minimizing precision-weighted prediction errors, where precision (i.e., estimated signal-to-noise in prediction errors) is encoded in the excitability of neuronal populations encoding prediction errors: c.f., Precision and Kalman gain in Bayesian filtering formulations of predictive coding.^1, 7, 8^ The empirical support for this scheme comes from studies showing that unpredictable stimuli (violating expectations) evoke increased gamma-band power and spiking that propagate feedforward, while predictable contexts enhance alpha/beta oscillations and connectivity in deep cortical layers, which can suppress gamma drive.^3, 4^

In Active Inference theory (AIF), predictive coding provides a computational foundation for how the brain continuously minimizes variational free energy through a hierarchical interplay of top-down predictions (priors) and bottom-up prediction errors (data)—evaluating prediction errors (perception), updating its model state and acting on the environment to fulfill these predictions (action).^9, 10^ Kolmogorov Theory (KT) formalizes this framework through Algorithmic Information Theory (AIT), where algorithmic *agents* optimize predictions by finding short (compressive) models of the world to optimize an Objective function. In the Bayesian framework of AIF, to achieve this, agents optimize the marginal likelihood of sensory input under a generative (i.e., compressive world) model of that input, where the log marginal likelihood or log evidence is bounded by variational free energy. Under the assumptions of predictive coding (i.e., continuous latent states and well-behaved random fluctuations), the objective function in AIF reduces to a mixture of precision-weighted prediction errors.^9, 10^ In KT, the Objective function represents the homeostatic and the tele-homeostatic (reproduction) goals of the agent, which indirectly translate into goals on minimizing prediction error^11–17^—see Figure 1. In both cases, measuring and managing prediction error is necessary.

**Figure 1:**
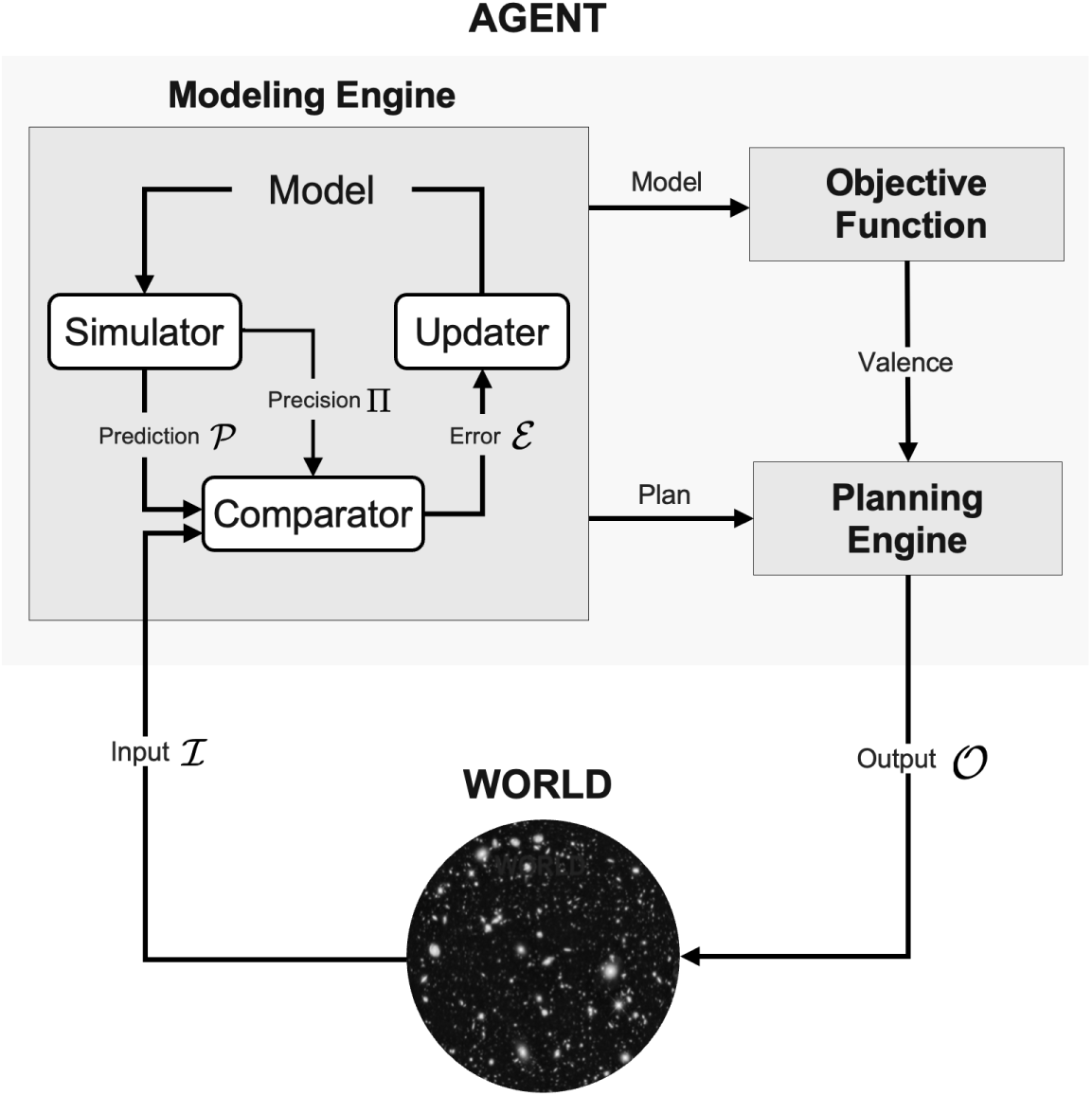
The Comparator in KT algorithmic agents. An algorithmic agent is an informationprocessing system with an Objective Function that interacts bidirectionally with the external world, creating and running compressive models, planning, and acting to maximize its Objective Function.^14–16^ The agent interacts dynamically with its environment, continuously exchanging data inputs (I) and outputs (O). The *Modeling Engine* formulates predictions (P) and the estimated precision (Π) of the prediction error to evaluate the precision-weighted prediction error (E) in the Comparator to update the Model. The *Comparator* is a key agent element monitoring the performance of the modeling engine and providing the information for model updates.

Although the predictive biological brain operates as a putative analog computational system, seemingly removed from the algorithmic digital agents described in KT, these paradigms are closely related. Both analog (e.g., brains) and digital algorithmic agents aim to model and track the world, a process that fundamentally relies on computing sensory prediction errors to update internal representations (information theory deals with both discrete and continuous notions of information under the same umbrella^18^). Predictive coding requires a *Comparator* mechanism to evaluate prediction error (E) to quantify how predictions P diverge from sensory input I to update predictions (see Figure 1). The Comparator assesses discrepancies, driving model adjustments to reduce E and aligning sensory and generative model dynamics. From a Bayesian filtering perspective, this can be read as Bayesian belief updating.^19, 20^

In this paper, we start by defining an information *encoding* framework inspired by amplitude modulation (AM) radio to explore, through neural mass modeling, the mechanisms by which prediction errors can be evaluated, identifying neural substrates and oscillatory instantiations of the Comparator function at the mesoscopic scale.

Our definition of the Comparator highlights essential distinctions between generating prediction errors and other important processes such as inhibition, gating,^21^ and routing,^22^ which can be read as predicting and instantiating the precision required to minimize precision-weighted prediction errors. In this setting, precision corresponds to an encoding of confidence, which plays a key role in weighting prediction errors that compete for belief updating. In Bayesian filtering schemes, this would correspond to optimizing the Kalman gain.^10^ In Psychology, this aspect of predictive coding has been associated with selective attention, attentional gain, and sensory attenuation. Physiologically, this context-sensitive aspect of predictive coding is usually attributed to cortical gain control or excitation-inhibition balance. The link between attention and physiological encoding of precision or uncertainty is nicely illustrated by the notion of communication through coherence.^3, 23–27^

In the binary encoding context, the Comparator is represented logically by the “exclusive” OR operation (XOR). XOR returns one if the inputs are different and zero otherwise, which can be expressed as E ∼ XOR(I, P) = I ⊕ P = (I ∧ ¬P) ∨ (¬P ∧ I), where E represents the prediction error, I the input and P the prediction. In predictive coding, the Comparator produces precision-weighted prediction errors that are crucial for model updating, directly informing adjustments to the generative model. Similarly, prediction errors are actively employed in Kalman filtering,^28^ enabling continuous updates to model parameters based on incoming sensory information.^19^ Thus, in the analog version, which is more appropriate for the physiological context, we generalize the Comparator function to evaluate the *weighted* difference between data and model, i.e.,

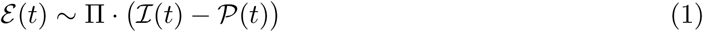

or some approximation to this quantity. Here, Π refers to the *Precision* (in reference to the Kalman filtering, see Appendix D) assigned to the prediction error, which we can think of as a multiplicative factor (or, more generally, an operator). Precision modulates the impact of the discrepancy between the model and data for model updates (loosely, the higher the precision, the greater the model update). More precisely, precision specifically refers to the inverse covariance of the prediction errors. However, in what follows, we will use the term somewhat loosely here to refer to inverse covariance (confidence) associated with a given signal (input, prediction, error).

### 1.2 The Role of Oscillations

There is a natural gradation from neuronal spiking to fast oscillations (gamma band), to slower rhythms such as beta, alpha, theta and delta.^29, 30^ These rhythms also exhibit distinct spatial coherence scales. Gamma oscillations tend to synchronize within local neuronal assemblies, whereas slower rhythms (e.g., theta, alpha) can coordinate across larger cortical networks.^29^ Consequently, information is encoded and integrated across these distinct spatiotemporal scales. Gamma oscillations (30–100 Hz) are critical for integrating information, feature binding, and encoding task-relevant signals across cortical circuits.^26, 29, 31^ In the motor cortex, gamma supports movement control and learning,^32^ while in the hippocampus it aids episodic memory formation.^33^ Disruptions in gamma are linked to neuropsychiatric disorders,^34, 35^ and gamma entrainment via sensory stimulation shows therapeutic promise.^36, 37^ Gamma synchronization—enabling communication through coherence—facilitates selective routing and feature binding.26, ^31, 38^

Animal studies further indicate that unpredicted stimuli evoke robust gamma (and theta) responses in superficial cortical layers, consistent with the feedforward transmission of prediction errors.^39^ Collectively, gamma rhythms are fundamental carriers of information for perception, attention, memory, and behavior.^40^ In contrast, alpha (8–12 Hz) and theta (4–8 Hz) oscillations are associated with inhibition, attentional gating, and cognitive control.^21, 41–45^ While gamma typically propagates feedforward, alpha is implicated in feedback processing.^46^ Moreover, alpha and beta oscillations have been shown to encode stimulus predictions and modulate spatial attention by selectively amplifying relevant signals.^47–50^

An important example of CFC^6^ is phase-amplitude coupling (PAC), where gamma oscillations are modulated by slower brain rhythms (theta, alpha, spindle, slow, and ultra-slow).^29, 51^ For example, theta oscillations organize gamma activity via PAC, with the theta phase modulating the gamma envelope to segment complex information (e.g., speech into syllabic units).^52–54^ Electroenchephalography (EEG), Magnetoencephalography (MEG), and Electrocorticography (ECoG) studies confirm that low-frequency responses track the temporal envelope of attended speech, while high-gamma envelopes selectively encode auditory information.^55–59^ Together, CFC among gamma, alpha, and theta rhythms establishes a dynamic balance between local processing and contextual coordination, which are hypothesized to subserve efficient and flexible neural computation.^60, 61^

### 1.3 Oscillatory and Predictive Coding alterations

#### 1.3.1 Disruption in AD and Other GABAergic Disorders

Alzheimer’s disease (AD) alters brain oscillations profoundly, shifting power toward slow frequencies (delta and theta) while reducing alpha, beta, and gamma activity.^62^ Gamma oscillations—central for precise timing and error signaling—decline with interneuron dysfunction, causing diminished gamma power and pathological hyper-synchrony in slower bands,^63^ as confirmed in human EEG/MEG during cognitive tasks and evoked potentials.^64^ These changes disrupt CFC, whereby gamma oscillations align with alpha/theta.^65–67^ AD patients show weaker theta/alpha to gamma PAC,^68, 69^ accompanied by reduced mismatch negativity (MMN) and delayed responses to unexpected stimuli,^70–72^ reflecting compromised predictive coding.^73^ Therapeutic strategies targeting gamma entrainment^74–76^ highlight the potential for restoring normal gamma oscillations and CFC patterns.

GABAergic dysfunction underlies these oscillatory and cognitive deficits,^77, 78^ particularly in parvalbumin-positive (PV) fast-spiking interneurons that enable fast oscillations. In prior work,^79^ we provided a laminar neural mass model simulating PV damage from amyloid-beta toxicity and pyramidal loss via tau pathology replicating AD’s full electrophysiological trajectory—from early hyper-excitability (increased alpha and transient gamma) to later hypoactivation dominated by slow waves—supporting the view that PV interneuron dysfunction drives these complex oscillatory dynamics. Here, we extend this modeling approach by linking these oscillatory phenomena to cognitive disruptions, i.e., to failures in predictive coding, high-lighting how impaired interneuron function leads to aberrant error signaling and a breakdown of hierarchical inference.

Similar PV-related impairments appear in autistic spectrum disorder (ASD), schizophrenia, and attention deficit hyperactivity disorder (ADHD): all show aberrant information processing and reduced sensory attenuation,^80–82^ alongside PV cell deficits.^83–86^ In ASD, PV neurons are fewer in several cortical areas,^83^ while morphological and functional PV alterations underlie gamma-band disturbances in schizophrenia and other conditions.^87^ Such interneuron dysfunction degrades high-frequency rhythms and disrupts predictive coding,^88–90^ undermining the top-down integration and error propagation that support normal cognition.

#### 1.3.2 Disruption by serotonergic psychedelics

Classical serotonergic psychedelics such as LSD, psilocybin, and DMT induce profound alterations in perception, cognition, and self-awareness by acting as 5-HT_2A_ receptor agonists. According to the REBUS model (“Relaxed Beliefs Under Psychedelics”), these substances reduce the precision of high-level priors, effectively weakening top-down constraints and allowing bottom-up signals to exert a stronger influence on conscious experience.^91^ Mechanistically, 5-HT_2A_ receptors densely populate layer V pyramidal cells, and their activation heightens neuronal excitability, disrupting the balance of cortical circuits that mediate weighted prediction errors.^92^ EEG and MEG studies show that this state is accompanied by broadband desynchronization across delta to gamma frequencies,^72, 92^ indicating reduced top-down control. In turn, electrophysiological measures of error processing are disrupted, reflecting an impaired ability to contextualize incoming stimuli.^71, 93^ Together, these findings support the view that psychedelics relax deep hierarchical priors, yielding both heightened error signals and a more “anarchic” brain state.^91^

Beyond spectral changes, empirical evidence from prediction error paradigms under psychedelics underscores the increase in error signals and impairment in pattern matching, as measured by oddball paradigms like the mismatch negativity (MMN) and the P300 component in eventrelated potentials (ERPs) in EEG/MEG.^94–96^ The P300, a positive ERP component associated with conscious evaluation of novel or target events and “updating” of context,^70, 71^ is reliably dampened by psychedelics. For instance, target oddball stimuli evoke a much smaller P300 amplitude compared to placebo.^72^ This reduction of P300 (and prolongation of its latency) implies that the brain’s ability to consciously evaluate and categorize events is affected—likely because relatively imprecise priors make it hard to decide what is contextually relevant. In a tactile MMN paradigm, psilocybin likewise reduced the neural responses to surprising stimuli (both in EEG and associated fMRI activity in frontal and sensory regions), indicating aberrant error processing in multi-sensory domains.^93^

Notably, some studies report that while “bottom-up” signals (like the basic MMN) remain, the later, more “top-down” components are disrupted.^93^ Using an auditory roving oddball (which assesses how MMN amplitude builds up with repeated standard stimuli), psilocybin was found to leave early error signaling largely intact: in one study, psilocybin did not significantly disrupt the MMN “memory trace” effect, whereas NMDA receptor blockade (ketamine) severely attenuated it.^71^ This suggests that pre-attentive error detection in the sensory cortex (which relies on short-term synaptic plasticity and local circuit NMDA function) is not abolished by 5HT2A agonism—consistent with the idea that psychedelics primarily affect higher-level inference rather than basic sensory change detection. This fits a picture in which psychedelics leave automatic error generation largely unimpeded or even enhanced, but interfere with the brain’s precision-weighting and integration of these errors into a coherent narrative.

In earlier work,^97^ we simulated the impact of serotonin 2A receptor activation on cortical dynamics. By modulating the excitability of layer 5 pyramidal neurons, our model reproduced the characteristic changes in EEG power spectra observed under psychedelics, including alpha power suppression and gamma power enhancement. These spectral shifts were shown to correlate strongly with the regional distribution of serotonin 2A receptors. Furthermore, simulated EEG revealed increased complexity and entropy, suggesting restored network function. The present work extends this research beyond the study of oscillatory power and frequency changes by focusing on the mechanisms for aberrant amplification of prediction error, which can be more directly linked with cognitive processes and (first-person) subjective experience.

### 1.4 The Laminar Neural Mass model (LaNMM)

Cortical function emerges from dynamics on cortical and subcortical hierarchies with multiscale architectural features.^98, 99^ These networks can be modeled using *connectomes*^100^ and node activity generated by physiologically realistic mesoscopic neural mass models (NMMs)^101^ to provide a high-level representation of the mean activity of large neuronal populations. A typical cortical column in the mammalian neocortex is estimated to contain on the order of 10^4^ neurons, with some regions or species exhibiting counts closer to 10^5^ neurons.^102^ For instance, in primary sensory cortices—such as the visual or somatosensory areas—a column is often characterized by roughly 10,000 neurons, although precise numbers can vary based on anatomical definitions and the specific cortical region. This mesoscopic scale—situated between single neurons and large-scale networks—provides a natural level of description for many electrophysiological phenomena. NMMs are designed to capture the aggregate dynamics of large neuronal populations by averaging individual activity into mean-field variables, such as average membrane potentials or firing rates.^103, 104^ Given that a cortical column comprises tens of thousands of neurons, the underlying assumption that microscopic fluctuations average out is largely justified. This averaging allows NMMs to reproduce key oscillatory and coupling phenomena, including cross-frequency interactions and the propagation of prediction errors, without modeling each neuron individually. Thus, NMMs are well-suited to describe mesoscopic dynamics within cortical columns, although they necessarily abstract away finer-scale microcircuital details.

We have previously introduced a *laminar* neural mass model (LaNMM)^105, 106^ optimized from laminar recordings from the macaque prefrontal cortex^107^ and designed to capture both superficial-layer fast and deeper-layer slow oscillations along with their interactions (cross-frequency coupling), such as PAC and amplitude-amplitude anti-correlation (AAC). The spectrolaminar motif is ubiquitous across the primate cortex.^108^ The LaNMM combines conduction physics with two NMMs—Jansen-Rit and Pyramidal Interneuronal Network Gamma (PING) subpopulations at slow and fast frequencies, respectively—to simulate depth-resolved electrophysiology.

Although the LaNMM is only one of several potential modeling frameworks, it provides a powerful and physiologically realistic tool to address the broader hypothesis that (1) information can be encoded both in ongoing signals and in their amplitude envelopes, and (2) prediction-error evaluation may be instantiated by cross-frequency coupling in cortical columns—particularly signal-to-envelope and envelope-to-envelope couplings. As we show, the LaNMM allows for a dynamic instantiation of the Comparator, facilitating the analysis of how prediction errors and top-down predictions may be integrated across different cortical lamina.

By featuring two interacting populations—each capable of autonomously generating oscillations—the LaNMM provides a physiologically grounded framework for studying cortical rhythms. In contrast, other models such as PING,^30^ Jansen–Rit (JR),^103^ Wendling,^109^ and Chehelcheraghi^110, 111^ differ in their structural assumptions and how they produce oscillations or crossfrequency coupling. Notably, the LaNMM does not rely on stochastic inputs to sustain its rhythms. For example, the Chehelcheraghi models require noise to induce inhibition-based gamma (ING) and CFC, whereas the LaNMM combines JR and PING sub-models, each generating slow or fast rhythms independently—leading naturally to phase-amplitude (PAC) and amplitude-amplitude coupling (AAC).

Furthermore, the LaNMM is explicitly designed with a cortical laminar architecture amenable to realistic physical measurements,^105, 112, 113^ distinguishing it from the lumped population models of PING, JR, and Wendling. While the PING model primarily produces gamma oscillations through an excitatory-inhibitory loop and the JR/Wendling models focus on alpha rhythms with additional fast inhibitory loops, the LaNMM naturally produces multiple frequency bands by leveraging laminar interactions. This makes it particularly suited for modeling observed laminar-specific oscillations, such as deep-layer alpha/beta modulating superficial-layer gamma, as seen in frontal and parietal cortices.^24^ Additionally, the model’s structure allows generalization beyond the alpha-gamma framework, accommodating different slow-fast frequency pairs (e.g., theta-beta, theta-gamma, beta-gamma) through parameter adjustments. This flexibility further enhances its relevance for modeling diverse cortical dynamics across brain regions (some examples are given in Appendix F).

Finally, the LaNMM fits squarely in the context of hierarchical processing and predictive coding. There exist evident asymmetries in hierarchical cortical connectivity, particularly in the distinct roles of feedforward and feedback pathways in sensory processing.^114, 115^ Anatomical studies have established that feedforward pathways typically originate from superficial layers and project to granular layers, while feedback pathways arise from deeper layers, connecting to non-granular areas, supporting a hierarchical processing model. Functionally, feedforward and feedback connections also differ in frequency domains. Feedforward signals often use gamma-band frequencies and are associated with prediction error transmission, while feedback signals utilize lower alpha and beta frequencies, hypothesized to convey predictions and in alignment with predictive coding models.^114^ In this framework, feedforward connections deliver prediction errors to higher-order areas, while feedback signals transmit top-down predictions to explain sensory input. Experimental studies support this model, showing that gamma coherence is stronger in feedforward pathways, while alpha/beta coherence is more prominent in feedback pathways, suggesting a mechanism where faster frequencies convey discrepancies and slower frequencies signal model-driven predictions.^114^

### 1.5 Amplitude Modulation for Information Encoding and Processing

In digital communication systems, information is typically represented as a sequence of binary digits. In contrast, analog systems and biological networks convey information via continuous signals. A classic example is amplitude-modulated (AM) radio, where a high-frequency carrier wave is modulated by a low-frequency signal so that the carrier’s amplitude (its envelope) encodes the desired information. Neural systems appear to operate on a similar principle, encoding information in aspects such as signal power, phase, or envelope.^116^ For instance, high gamma-band activity in the auditory cortex exhibits amplitude modulation that closely tracks the stimulus envelope,^117^ and Pasley et al. (2012)^118^ demonstrated that speech could be reconstructed from the high gamma power envelope extracted from ECoG recordings.

In AM radio, information—such as speech or music—is encoded as variations in the dynamic amplitude (envelope) of a high-frequency carrier wave. In more detail, the fast carrier signal with frequency *f_c_* and amplitude *A_c_*, expressed as *C*(*t*) = *A_c_* cos(2*πf_c_t*), is modulated by a slower base-band (information) signal *m*(*t*) to produce *S*(*t*) = *A_c_* [1 + *a m*(*t*)] cos(2*πf_c_t*), where *a <* 1 is the *modulation index* ensuring the envelope reliably follows *m*(*t*). At the receiver, demodulation extracts the original information from the envelope variations of *S*(*t*).^119^

In our framework for error evaluation, and as further discussed in Ruffini et al. (2025),^120^ we assume that oscillatory information is encoded in the slow (top-down) *signal*, its *envelope*, or in the fast signal’s *envelope*. For this reason, rather than PAC, we will focus on signal-to-envelope coupling (SEC). As the dynamic impact of error signals is proportional to their power, “loud” (i.e., precise) signals exert a stronger effect on predictive processing (i.e., belief updating in higher hierarchical levels). In this view, the Comparator mechanism at each hierarchical level does not operate on the full detail of the gamma signal but rather on a coarse-grained version— the amplitude envelope—that approximates the difference between internal predictions and incoming sensory data. A successful match yields a lower power gamma signal with a reduced modulation index, whereas a discrepancy results in a modulated signal conveying the prediction error.

As an extension of this framework, the Hierarchical Amplitude Modulation (HAM) model^120^ proposes that information is encoded not only in the raw oscillatory signals but also in their successively extracted envelopes, establishing a multiscale coding scheme that reflects the brain’s hierarchical organization—and implicit depth of its generative world model.

### 1.6 Paper Roadmap

In the following sections, we detail how our model capitalizes on the above encoding mechanisms. First, we show that the LaNMM provides concrete mechanisms for SEC and EEC. In SEC, slow-wave activity modulates the amplitude envelope of fast oscillations (analogous to phase-amplitude coupling), while in EEC, slower envelopes modulate amplitude-amplitude interactions, thereby adding an extra layer or order of predictive control. In brief, we implicitly associate (fast) phase amplitude coupling with the encoding of first-order content of prediction errors, while the (slow) amplitude-amplitude coupling is associated with the encoding of secondorder context in which content is inferred, where the context can be regarded as the precision (i.e., inverse variance) or second-order statistics of first-order prediction errors.^8, 121^

Next, we demonstrate how the LaNMM can provide weighted prediction-error evaluation, i.e., computing an approximate difference between incoming sensory data (as captured in input signal envelopes) and internal model predictions (either as signals or their envelopes). These mechanisms enable the Comparator to operate across multiple levels and timescales—generating fast prediction-error signals via SEC and slower context-sensitive weighted gating signals via EEC.

Finally, to explore how alterations in the LaNMM and prediction error function can shed light on the adverse effects observed in certain pathological or altered states, we discuss how deficits in fast inhibitory synapses, as seen in Alzheimer’s Disease, and increased glutamate receptor gain, characteristic of serotonergic psychedelic states, may disrupt the Comparator process and compromise effective information processing.

## 2 Methods

Figure 3 provides an overview of the Comparator mechanism in predictive coding, which we will implement in our LaNMM. Here, incoming “input” I is combined with a “prediction” P, producing a precision-weighted “error” signal E propagating up the hierarchy.

Information is encoded in the “fast envelope” of the input signal I and in the prediction signal P itself, while input precision (gain) is encoded in the slow envelope of the input (envelope of the envelope, i.e., average power) and prediction precision in the envelope of the prediction signal. More specifically, the envelope of the prediction signal encodes the precision of the prediction Π_P_ and the inverse precision of the prediction error signal (i.e., covariance or uncertainty), while the envelopes of input and error encode their precision, Π_I_, and Π_E_ respectively (inverse covariance or confidence).

In line with predictive coding frameworks, we aim to show how the LaNMM can explicitly encode mismatch signals (errors) that update and refine internal predictions, thereby emulating a biologically plausible Comparator process at multiple cortical levels. In this paper, we will focus on local processing at a single column and the implementation of modulation, demodulation, and Comparator mechanisms. The analysis of coupled columns for a full implementation of predictive coding is left for future work.

For the implementation of the Comparator, we rely on oscillations and their cross-frequency coupling properties. In our demonstration, we focus on a particular combination of inputs. However, the mean inputs of the LaNMM nodes in a whole-brain model will vary as a function of their connectivity to other nodes. To illustrate the range of applicability of the model, we start by characterizing the dynamics and CFC properties of the model in the input landscape.

### 2.1 Dynamics and Cross-Frequency Coupling in the LaNMM

To analyze the dynamics of the model—prior to our implementation of the Comparator function— we first obtained the bifurcation diagram of the model as a function of mean inputs to the two pyramidal populations. Figure 2 presents the two-parameter bifurcation diagram of the model for (*µ_P_*_1_ *, µ_P_*_2_ ), obtained using the continuation software AUTO-07p.^122^ Other model parameters were set as in Sanchez-Todo et al. (2023).^105^ Starting at *µ_P_*_1_ = 0, the system exhibits a stable fixed point and evolves toward a steady state. Along the *x*-axis for *µ_P_*_2_ = 0, the system undergoes a saddle-node on an invariant cycle (SNIC) bifurcation around *µ_P_*_1_ ≈ 104 (depicted in green), after which it transitions to periodic dynamics. Increasing *µ_P_*_1_ further leads to two Fold of Limit Cycles (FLC) bifurcations, represented by the dashed red lines. Finally, around *µ_P_*_1_ ≈ 364 the periodic dynamics disappear at a supercritical Hopf bifurcation HB^+^ (purple).

**Figure 2:**
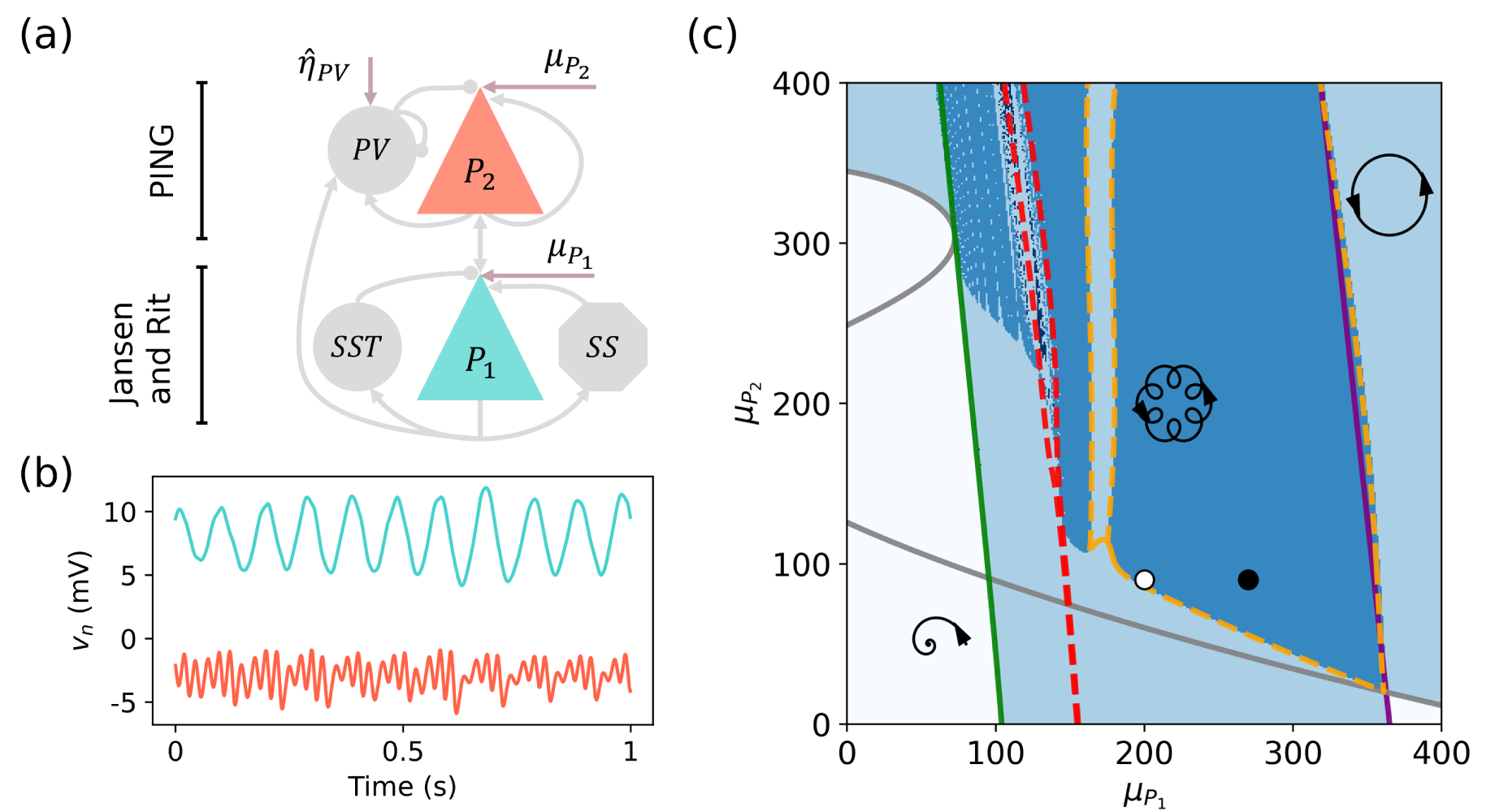
Laminar Neural Mass Model Overview. (a) Laminar neural mass model architecture. Driving inputs studied in the model (all glutamatergic) are displayed (*µ_P_*_1_ is the mean input parameter to *P*_1_, *µ_P_* _2_ is the mean input parameter to *P*_2_, as well as stochastic zero mean input to PV). (b) Sample outputs of the LaNMM model (membrane potential perturbation of *P*_1_ and *P*_2_ populations). (c) Two-parameter bifurcation diagram of the LaNMM model as a function of the external inputs *µ_P_*_1_ and *µ_P_*_2_ . Symbols indicate the dynamical regimes within the colored regions: steady-state (white), periodic (light blue), and quasi-periodic (dark blue). The lines represent bifurcations, as described in the main text. The white dot marks the mean input parameter values used in Sanchez-Todo et al. (2023),^105^ (*µ_P_*_1_ *, µ_P_*_2_ ) = (200, 90) Hz. The black dot indicates the representative values chosen in this study, (*µ_P_*_1_ *, µ_P_*_2_ ) = (270, 90) Hz, which place the model deeper into the quasi-periodic regime and display a stronger EEC (see Figures 4 and G2).

**Figure 3:**
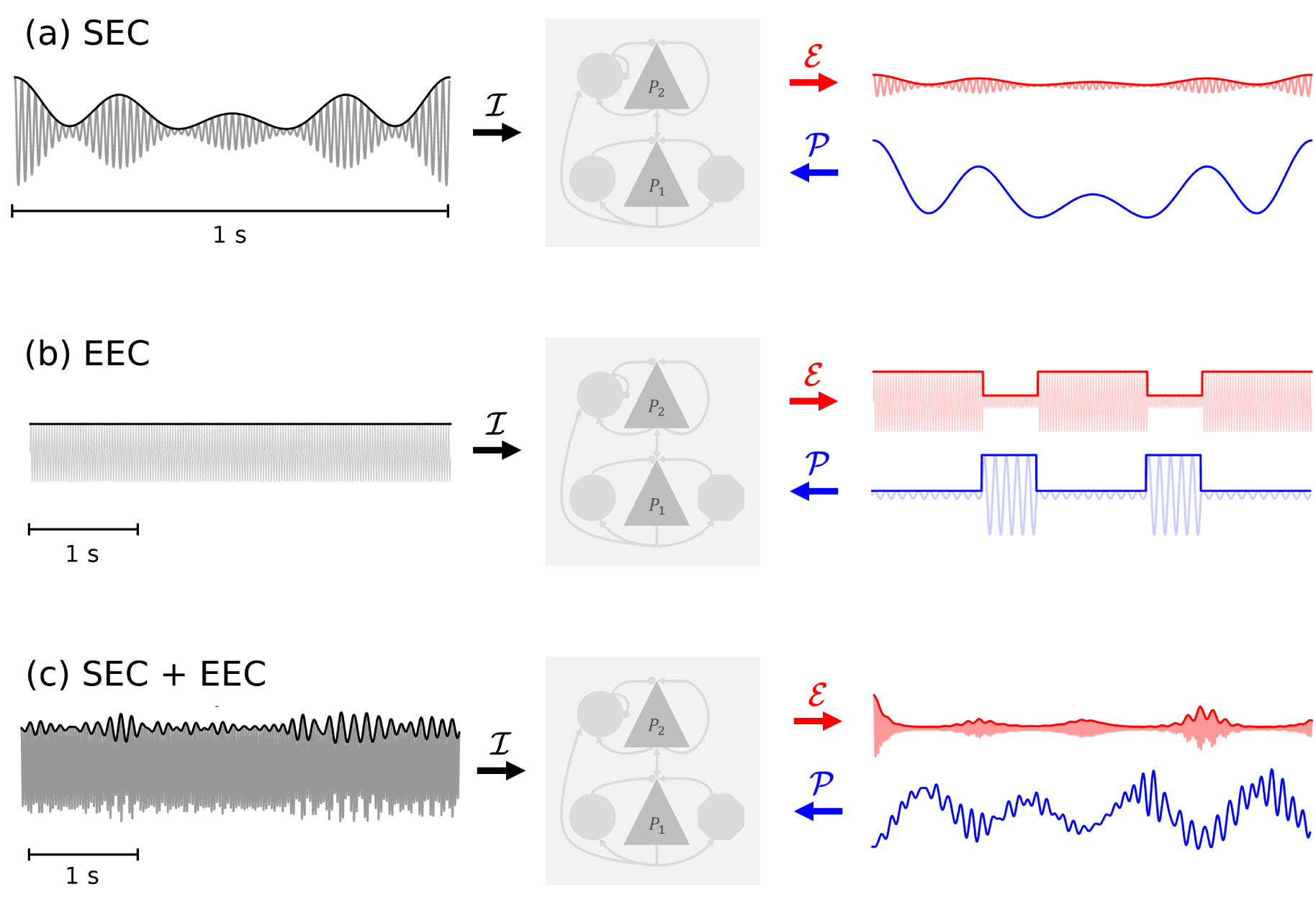
Laminar model concept implementation of the evaluation of prediction error (E) via two timescales and two mechanisms: a) Signal-Envelope coupling (SEC) mediated evaluation of prediction error, where the incoming information is encoded in the (fast) envelope of the carrier, and the prediction is directly the slower signal; b) Envelope-Envelope coupling (EEC) implementation of precision computation (digital simplification), where the precision of prediction errors is modulated by the envelope of the slower signal flowing downstream. Panel (c) illustrates the combination of both mechanisms, where precision (very slow envelope) and predictions (signals) are encoded into the same signal (P). The LaNMM (center) includes two bidirectionally interacting subpopulations.^105^ Bottom-up input data (I) arrives at the PING subpopulation pyramidal cell *P*_2_ encoded in the fast oscillation envelope, while top-down weighted predictions (P) arrive encoded in slower signals (alpha band in the model) at the bottom JR population pyramidal cell *P*_2_ or at the PV cell in the PING subpopulation. The interaction alters the dynamics of *P*_2_, which produces an approximate, weighted difference of input and prediction E encoded in the envelope.

This bifurcation scenario closely resembles that of the coupled Jansen-Rit model discussed in Clusella et al. (2023).^123^ Notably, without external inputs to *P*_2_, the system is unable to generate gamma rhythms within this range. While the overall structure remains consistent as *µ_P_*_2_ increases, an additional supercritical Hopf bifurcation HB^+^ (gray) emerges. Of particular interest is the region where two limit cycles coexist, leading to a torus bifurcation (TR) and subsequently driving the system into a quasi-periodic regime. The white dot marks the mean input parameter values used in Sanchez-Todo et al. (2023),^105^ (*µ_P_*_1_ *, µ_P_*_2_ ) = (200, 90) Hz. The black dot indicates the values chosen for this study, (*µ_P_*_1_ *, µ_P_*_2_ ) = (270, 90) Hz, used in recent modeling work on psychedelics and AD.^97^

Next, we analyzed the cross-frequency properties of the model. *Signal-envelope coupling (SEC)*— which is related but not equivalent to PAC^107, 124–128^ — can be used to implement the error evaluation/comparison process E at relatively fast time scales (dictated by the low-frequency signal). Here, a slower oscillatory rhythm (encoding predictions) modulates the envelope (or the overall magnitude) of faster oscillations (encoding the incoming sensory data) at the slower timescale. This mechanism allows the slower predictive oscillation to interact with information encoded in the fast oscillation at a “coarse-grained” level, where discrepancies are identified. Both the phase and the instantaneous value of the slower oscillation impact this modulation, aligning the processing of fast inputs with the slower rhythm and enabling the system to compute the necessary error signal.

*Envelope-Envelope Coupling (EEC)*, sometimes referred to as Amplitude-Amplitude Coupling (AAC) or power coupling, can act as a suppression or *gating mechanism at a slower timescale*, which acts by modulating the envelope of faster oscillations according to the envelope of a slower rhythm. This coupling creates windows of increased or decreased excitability, effectively allowing or restricting the propagation of error signals based on the slow oscillatory envelope. Through this gating function, EEC facilitates selective routing by maintaining or suppressing the transmission of specific error signals, extending routing effects over broader timescales. This mechanism we associate with the modulation of prediction error signaling to effectively broadcast precision-weighted prediction errors. EEC allows predictive coding circuits to remain dynamically sensitive to shifts in attentional or contextual relevance. By sustaining periods of high or low excitability based on slower rhythmic inputs, EEC supports long-term gating of information flow, ensuring that predictive models align with high-level task demands or goals (e.g., attention, vigilance). However, EEC can also be used as a graded inhibitory mechanism capable of encoding contextual information to be contrasted with the input power at longer timescales. From a statistical perspective, the greater integration time is mandated by the need to accumulate evidence for the precision of prediction errors. This follows because the estimation of precision requires the pooling of instantaneous prediction errors over time (in the same way that a statistician would pool prediction errors over multiple observations to estimate the standard error of her estimated effect size).

The LaNMM model as described in Sanchez-Todo et al. (2023)^105^ was used to investigate the interactions between the two pyramidal populations in the neural mass and the SEC and EEC mechanisms under varying external inputs *φ_e_*_1_(*t*) and *φ_e_*_2_(*t*). Simulations were conducted across a two-dimensional parameter space by performing a sweep over the mean input values *µ_P_*_1_ and *µ_P_*_2_ over a range from 50 to 400 in steps of 5 Hz. For each parameter pair, we integrated the LaNMM model using a fixed time step (Δ*t* = 1 ms) and a total simulation time of ∼90 s, discarding an initial transient. Depending on the desired drive condition, the model was run in one of three modes: *P*_1_ drive, *P*_2_ drive, or PV drive. External inputs were generated using a multiscale noise generator that produces a slow random-walk drift and a fast white-noise component (see Appendix G), which were injected into the appropriate population (either *P*_1_, *P*_2_ or, via an extra glutamatergic synapse, PV).

SEC was quantified by computing the Pearson correlation coefficient between the band-pass alpha band signal of *P*_1_ and the amplitude envelope of *P*_2_’s gamma signal. EEC was similarly assessed by correlating the amplitude envelopes of *P*_1_ in the alpha band (8 − 12 Hz) and *P*_2_ in the gamma band (30 − 50 Hz). In addition, we computed the power in the alpha and gamma band in both pyramidal populations and the Population Excitation-Inhibition Index (PEIX) to characterize the sigmoid saturation of fast (*P*_2_) and slow (*P*_1_) oscillatory activity in the model to complement the analysis (see Appendix A for details).

These calculations were performed for the standard LaNMM parameters (other than inputs), but also for the scenarios of PV dysfunction, psychedelics, and their combination (see Appendix H)

### 2.2 Comparator I: Implementation Using SEC

#### 2.2.1 Error suppression

We can formulate a first hypothesis, regarding the implementation of the comparator with the LaNMM: for a given input, if the correct prediction is applied (i.e., an alpha signal matching the gamma envelope of the input signal), the output error should be low, which can be measured using the power of the *P*_2_ output envelope—see Figure 3.

To demonstrate this, we created synthetic signals to be used as input and prediction messages. The input signal was generated by amplitude modulating a 40 Hz carrier signal with an alpha-band (8–12 Hz) envelope extracted from filtered white noise. To ensure non-zero values, a constant offset of 0.2 mV was added to the envelope and the carrier wave. The prediction signal was designed to either match this alpha envelope (match condition), remain constant (flat condition), or originate from an independent alpha source (wrong condition). Input and prediction signals can be visualized in Figure 5a. These conditions allowed us to investigate the role of input-prediction alignment in modulating the network’s response.

Given that *P*_1_ is an excitatory population and that the SEC for *P*_1_-driving inputs is positive (see Figure 4), we expected—and confirmed—that for effective suppression of the input signal in the match condition, the prediction signal had to be phase-shifted by 180° (i.e., inverted). This inversion allowed the prediction to counteract the excitatory input effectively. In contrast, for the PV population, which is inhibitory and has positive SEC for PV-driving inputs, the prediction had to be applied in phase with the input signal to achieve suppression. Further details on the variation of power suppression depending on the phase of the matching prediction are shown in Appendix C.

**Figure 4:**
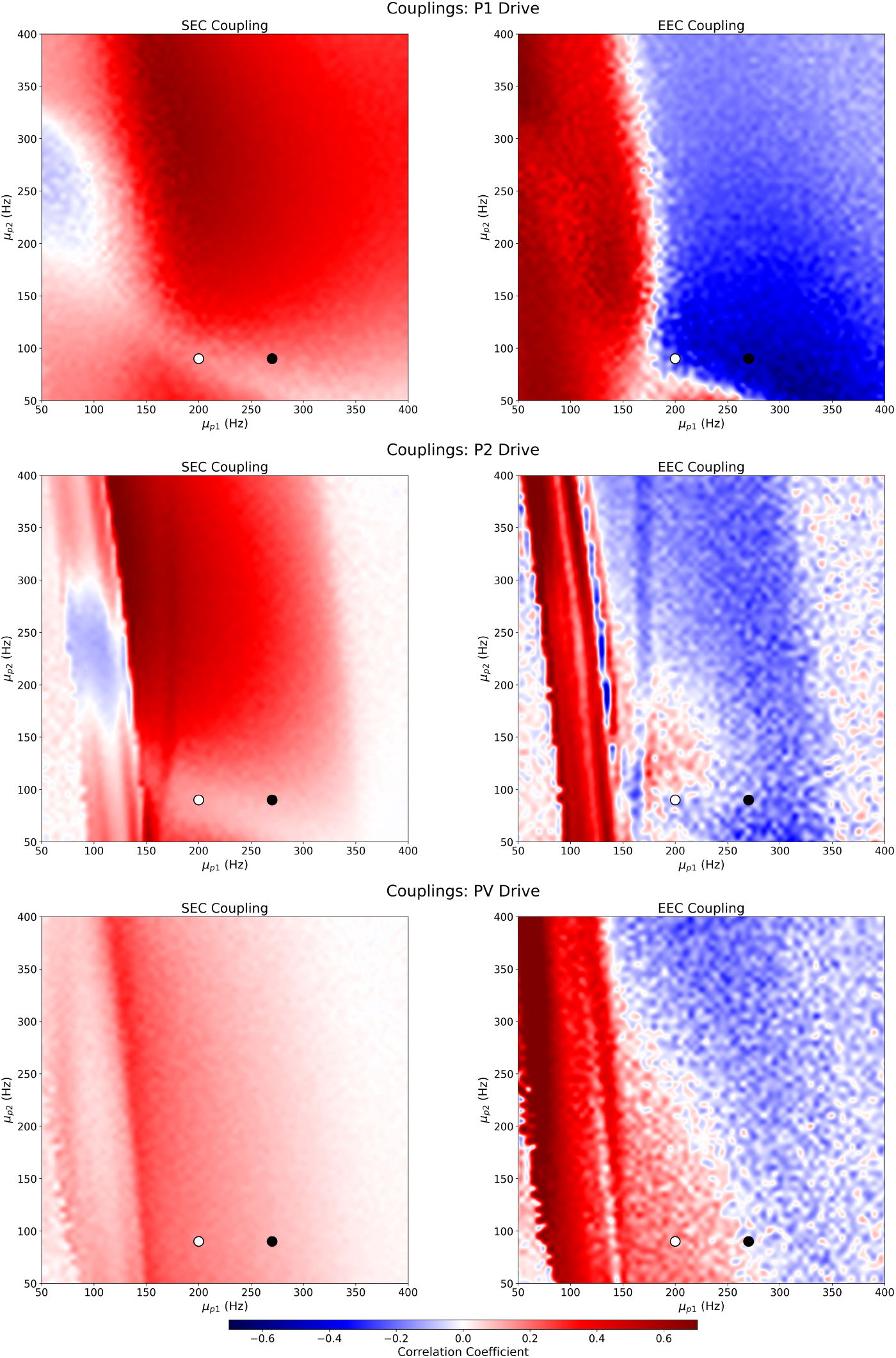
SEC (left column) and EEC (right) between neural populations *P*_1_ and *P*_2_ for random inputs to *P*_1_ (top row), *P*_2_ (middle), and PV (bottom). Both *P*_1_ and *P*_2_ receive a constant mean external drive (*x* and *y* axis), with zero mean noise injected into one of them via a glutamatergic synapse. This dual-input strategy allows the model to capture both the steady excitatory drive and the stochastic fluctuations affecting the desired population. The left column depicts the correlation between the alpha signal of *P*_1_ and the gamma envelope of *P*_2_ (SEC), while the right column displays the correlation between the alpha envelope of *P*_1_ and the gamma envelope of *P*_2_ (EEC). The white dot marks the mean input parameter values used in Sanchez-Todo el al. (2023),^105^ (*µ_P_*_1_ *, µ_P_*_12_) = (200, 90) Hz. The black dot indicates the values chosen for this study, (*µ_P_*_1_ *, µ_P_*_2_ ) = (270, 90) Hz.

**Figure 5:**
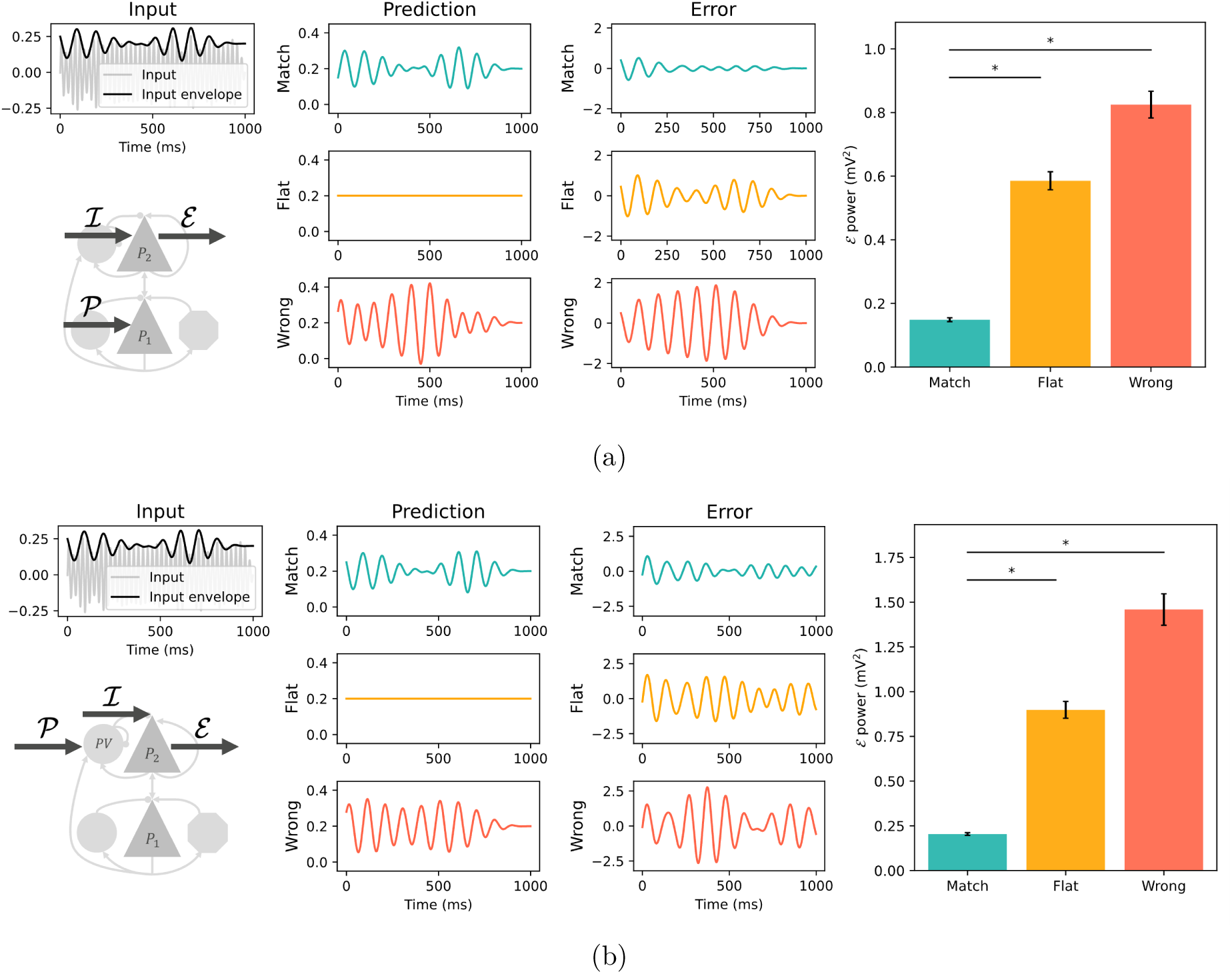
SEC suppression scheme, prediction signal sent to *P*_1_ (a) or PV (b). If the prediction signal matches the input (blue traces), the error signal—*P*_2_ output envelope—is suppressed, as shown by the reduced power (right bar plot). We compare the case of a matching prediction with the case of a flat prediction signal (orange traces) and a wrong prediction signal (red traces). The asterisk denotes a significant difference over multiple simulations in a paired t-test (p *<* 1e^−5^), a convention followed in all subsequent figures.

These signals were then used to drive the LaNMM. The generated signals were applied as inputs, modifying the membrane potential perturbation *u_n_* of the corresponding populations. The inputs and predictions were scaled by gain factors *G*_I_*, G*_P_. Simulations were run for a total duration of 100 seconds with a sampling frequency of 1000 Hz.

After running the model, the membrane potential perturbation of the *P*_2_ population, *u_P_* _2_, was extracted and analyzed. This output signal was band-pass-filtered between 30 and 50 Hz to isolate gamma activity, and its envelope was computed using the Hilbert transform. The envelope *P*_2_ signal was then further filtered in the alpha-band (8–12 Hz) to extract the error signal (E). Finally, the power of this alpha-band signal was computed as a measure of error power, allowing us to quantify the impact of different predictions on the suppression of mismatch signals.

We first optimized the gain factors *G*_I_*, G*_P_ to determine the conditions that resulted in maximal suppression of the error signal in the “match” condition. This optimization was performed by varying the gains and evaluating the resulting error power (see Appendix B).

Using these optimized parameters, we ran ten simulations using different seeds to generate the signals and compared the power of the error signal across the different prediction conditions. This analysis allowed us to assess the extent to which the alignment between the prediction and the input influenced the suppression of mismatch-related activity within the model.

#### 2.2.2 Computation of prediction error

The above study is a special case of a more general statement: for a given input signal encoding a message (in the gamma envelope) and a given prediction signal encoding a different message (as an alpha band oscillation), the output signal (encoded as the gamma envelope of the *P*_2_ output) will approximate the difference of the signals, i.e., a prediction error approximation.

The hypothesis can be formalized as E = *a* · I + *b* · P + *η*^, where E is the *P*_2_ output envelope (and prediction error estimate) produced by the LaNMM, as explained in the previous section, I is the input message (fast oscillation envelope), P is the prediction message (slow oscillation signal), and *η*^ denotes unmodeled noise. Here, *a* and *b* are constants (real numbers) that scale the input and prediction, respectively, and we expect *a* and *b* to be of opposite sign (subtraction).

### To determine *a* and *b*, we optimize the loss function

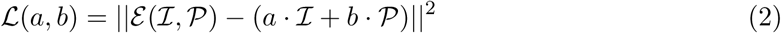

Using this loss function, we find *a* and *b* with which the LaNMM output is best approximated as a linear function of input and prediction.

To demonstrate that the minimal loss is meaningful, we define an alternative loss function,

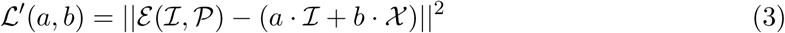

where X denotes a random assignment of mismatched predictions. We can then verify that, for a given pair (I, P) used as input and prediction to the LaNMM, the minimization of L results in a lower loss than the minimization of L^′^. We assume that the unrelated predictions X are instantiations of different alpha signals with the same mean and amplitude as the prediction sent to the LaNMM, obtained with different seeds.

To generate information-rich output and subtraction messages and avoid power suppression of the input, the signals used as predictions did not match the input signals (i.e., they were generated with different seeds than the input, corresponding to the “wrong” prediction case in the previous section).

We generated ten such pairs of (I, P) and ten unrelated predictions X for each pair, and minimized L and L^′^ in each case. To understand whether the error computed with the LaNMM was best approximated by a linear combination of (I, P), we ran an independent samples t-test comparing the distribution of L and L^′^values. We carried out this study for the prediction sent to the *P*_1_ population and for the prediction sent to the PV population.

### 2.3 Comparator II: The role of EEC for precision weighting

Here, we study the idea that EEC (anticorrelation) can be used to modulate the precision of prediction errors within the column—see Figure 3. We assume that the precision of prediction errors, Π_E_ , is encoded in their lower-frequency envelope (longer timescale averaged power) and show how EEC can be used to amplify or attenuate the precision of error by applying topdown precision-modulation signals Π_P_. Implicitly, we assume that the confidence in inputs (the precision of the incoming error from bottom-up layers), Π_I_, is encoded in the amplitude of their envelopes, too. Given the low EEC coupling with PV drive (Figure 4), we did not expect the error computation to be successful with EEC, when the precision-modulating signal was applied to the PV population, and we did not simulate this scenario.

As input signal, we used a non-modulated 40 Hz carrier signal with mean 2 mV and amplitude 2 mV, to ensure that the input was always positive. To obtain a precision-modulation signal, we created a random square wave of 100 seconds by adding 5 random steps of duration 10 seconds and amplitude 1 mV to a flat signal of 1 mV. Such a square wave represents the input precision Π_I_. We amplitude-modulated a 10 Hz carrier signal to obtain the precision-modulation signal. We added a constant offset of 2 mV to the resulting signal to ensure non-zero values. To evaluate the effect of the precision-modulation signal, we used as a contrast condition a “flat” precision-modulation signal (no modulation) oscillating at 10 Hz (with mean 2 mV and 2 mV amplitude). Input and prediction-modulation signals can be visualized in Figure 6. As before, the input and precision-modulating signals were scaled by gain factors *G*_I_ = 1*, G*_P_ = 10, which were optimized to result in maximal modulation of the error precision (such optimization is detailed in Appendix B).

**Figure 6:**
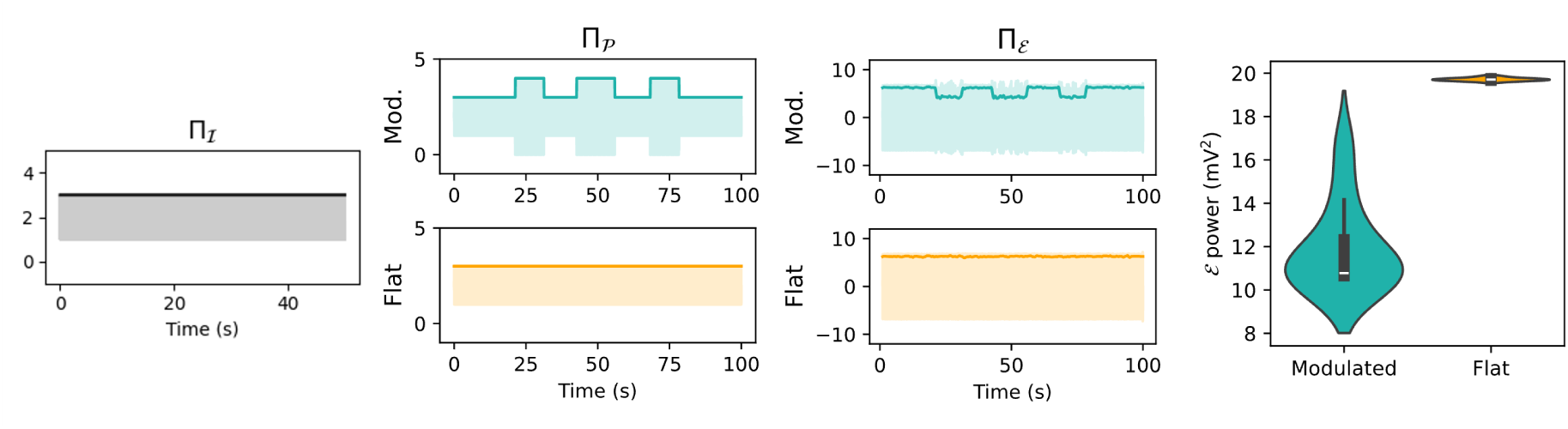
EEC suppression scheme. The precision modulation signal presented to *P*_1_, Π_P_, successfully attenuates the error precision Π_E_ (*P*_2_ output slow envelope). The error precision is reduced by the precision-modulation signal, as shown by the reduced error power (right violin plot). We compare the case of a precision-modulation signal (blue traces) with the case of a flat precision-modulation signal (orange traces, no modulation).

After running the model, the membrane potential of the *P*_2_ population was extracted and analyzed. The output signal was band-pass-filtered between 30 and 50 Hz to isolate gamma activity, and its envelope was computed using the Hilbert transform. The envelope *P*_2_ signal was then further low-pass filtered below 1 Hz to extract the error signal precision (Π_E_ ). Finally, the gamma power of the *P*_2_ output was computed and compared for the precision-modulation and flat modulation signals as a measure of error attenuation. We ran ten simulations using different seeds to generate the signals and evaluated the difference across conditions using paired t-tests.

#### 2.3.1 Modulation of precision

Here we aim to show that for a given input signal encoding a message with some precision Π_I_ (in the slow envelope of gamma) and a given prior signal encoding some precision modulation Π_P_ (in the alpha signal envelope), the output signal will encode an approximate subtraction of the respective precisions, reflected in the slow envelope of the *P*_2_ output.

In this case, we used as input messages signals with varying precision Π_I_, obtained in the same way as the precision-modulating signals described in the previous section. Prior messages encoding the precision modulation were obtained in the same way.

We observed that such simple input and prior messages of amplitude 1 mV, the normalized *P*_2_ output slow envelope (between -1 and 1), Π^^^_E_ , approximates the direct subtraction of input and prior precisions well. We thus computed directly the MSE between the slow envelope of the *P*_2_ output and the subtraction of input and prediction precisions (long time scale envelopes) as a measure of correct precision modulation:

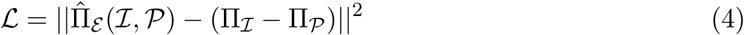

We will refer to this as the *consistent* case since the input and prediction provided to the LaNMM are the ones used in the subtraction to compute the MSE.

To demonstrate that such MSE obtained is meaningful and that the modulation computed by the LaNMM approximates the subtraction of input and prior precision well, we can also compute the MSE between the LaNMM output slow envelope and the subtraction of input and an unrelated prior precision Π_X_ (inconsistent case),

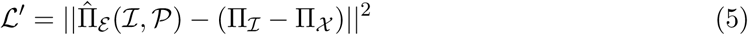

The unrelated prior precision X is constructed in the same way as the prior sent to the LaNMM but with another seed to define the random signal generation.

As in the SEC case, the signals used as priors did not match the input precision signals (i.e., they were generated with different seeds than the input, corresponding to the “wrong” prior case in the previous section). We generated 10 pairs of (I, P) to be used as input and prior to the LaNMM and one unrelated prior X for each pair, and computed L and L^′^. We ran a paired t-test comparing the distribution of MSE values for the consistent and inconsistent cases.

### 2.4 Disruption of Comparator Function

#### 2.4.1 Serotonergic Psychedelics

We then studied disruption of the comparator function due to the interaction with classical serotonergic psychedelics (*psychedelics* hereafter). Based on previous modeling work,^97^ the increased excitability to glutamate inputs observed under psychedelics was implemented in the model as an increase of the average synaptic gain of excitatory connections to the *P*_1_ population, *A_P_*_1_ . Three glutamatergic connections into *P*_1_ are present in the LaNMM: connections from the *SS* and *P*_2_ populations to *P*_1_, and external noise perturbation *e*_1_ to *P*_1_ (see Figure 2). See also the empirical changes in excitability estimated using dynamic causal modelling in (Muthukumaraswamy et al., 2013).^92^

We first analyzed the alterations induced by psychedelics in the oscillatory behavior of the LaNMM by computing the alpha and gamma power of the *P*_1_ and *P*_2_ populations for different values of the *A_P_*_1_ parameter. Based on those results, we chose a value of *A*^Ψ^ = 4.5 mV to represent the acute phase of psychedelics. For this value, the alpha power in the *P*_1_ population was strongly reduced in the model, while the gamma power in the *P*_2_ population was substantially increased.

We analyzed the effects of these alterations in the LaNMM computation of prediction error using the SEC and EEC mechanisms. We first computed the power of the error signal (*P*_2_ output gamma envelope) obtained in the SEC scenario in the *baseline* condition (*A*^0^ = 3.25 mV, default value in the LaNMM and in Jansen-Rit^97, 103, 105^) and the *psychedelics* condition, in the case where a prediction signal matching the input signal was applied.

We then studied the alterations in the EEC mechanism for precision computation. As in the SEC case, we computed the gamma power of the error signal (*P*_2_ output slow envelope) in the baseline condition and the psychedelics condition, in the case where a 10 Hz precision-modulating signal was applied to *P*_1_.

#### 2.4.2 PV disinhibition in Alzheimer’s Disease

The inhibitory deficits associated with AD were implemented in the model as a reduction in the strength of inhibitory synaptic connections from the PV population to the *P*_2_ population, *C_P V_* _→*P* 2_, based on previous computational models of AD-related dysfunction.^79^

We first analyzed the alterations induced by AD in the oscillatory behavior of the LaNMM by computing the alpha and gamma power of the *P*_1_ and *P*_2_ populations for different levels of PV dysfunction. Based on the work in Sanchez-Todo et al. (2024),^79^ we selected a level of inhibition reduction for different stages of disease progression: healthy (baseline value, *C_P V_* _→_*_P_* _2_ = 550), Mild Cognitive Impairment (MCI) (*C_P V_* _→_*_P_* _2_ = 300) and AD (*C_P V_* _→_*_P_* _2_ = 140).

We first evaluated the alpha and gamma power alterations in the different stages of AD due to variations in the *C_P V_* _→_*_P_* _2_ parameter. We then analyzed the effects of these alterations in the LaNMM computation of prediction error using the SEC and EEC mechanisms.

We computed the power of the error signal (*P*_2_ output gamma envelope) obtained in the SEC scenario for each AD stage, using a prediction signal matching the input. We then studied the alterations in error precision computation using EEC. To do so, we used a flat input oscillating at 40 Hz and a prior signal encoding some precision-modulation oscillating at 10 Hz. We computed the power of the error signal in each AD stage as a measure of the EEC error suppression by the precision-modulating signal.

## 3 Results

### 3.1 The LaNMM Sustains SEC and EEC for a Wide Range of Mean Inputs

We began by examining the behavior of the system when random inputs were presented to *P*_1_, *P*_2_, and PV. Figure 4 displays maps indicating the mean positive SEC and negative EEC values. These results reveal that in a wide range of configurations, the system naturally yields the desired combination of positive SEC and negative EEC. This provides a baseline for how the laminar model processes random inputs before introducing comparator mechanisms.

The results of the simulation of SEC and EEC highlight the robustness of EEC and SEC couplings across wide ranges of the parameter space. As a result of robust EEC with *P*_1_ driving (Figure 4), we see that envelope coupling provides a mechanism for suppression of competitive oscillations. For example, alpha power (evaluated by its envelope) can be used to suppress the gamma envelope or vice versa. However, the same applies in the other direction: fast oscillation power can be used to regulate slow oscillation activity. This mechanism can subserve gating by inhibition and predictive routing, as is further discussed below. Figures A1 and A2 in the Appendix display maps of the computed power within the alpha and gamma bands for both *P*_1_ and *P*_2_, revealing how power distributions vary with changes in the external drives *µ_P_*_1_ , *µ_P_*_2_ and *µ_P V_* .

### 3.2 SEC subserves Fast-Time Comparator Function

We first employed the SEC mechanism to implement the Comparator function (i.e., the difference between input and prediction). Two primary configurations were tested: providing the prediction to *P*_1_ and providing the prediction to PV. Figure 5a illustrates the suppression of error signal power in the first case. The second configuration (prediction to PV) is shown in Figure 5b, demonstrating that both achieve a similar comparator function using SEC. Indeed, in both cases, sending a prediction signal matching the input signal results in lower prediction error (measured as the E power) than sending a flat prediction or a wrong prediction.

Figure 7 illustrates the computation of prediction error when the prediction is provided to the *P*_1_ population. The main results for the PV configuration are also shown. As shown in Figure 7d, the loss value with the original prediction is always lower than the loss value with alternative predictions. This demonstrates that the linear combination of the input and prediction signals sent to the LaNMM explains the LaNMM output better than the combination of the input with other arbitrary prediction signals (i.e., that L^∗^ is minimal).

**Figure 7:**
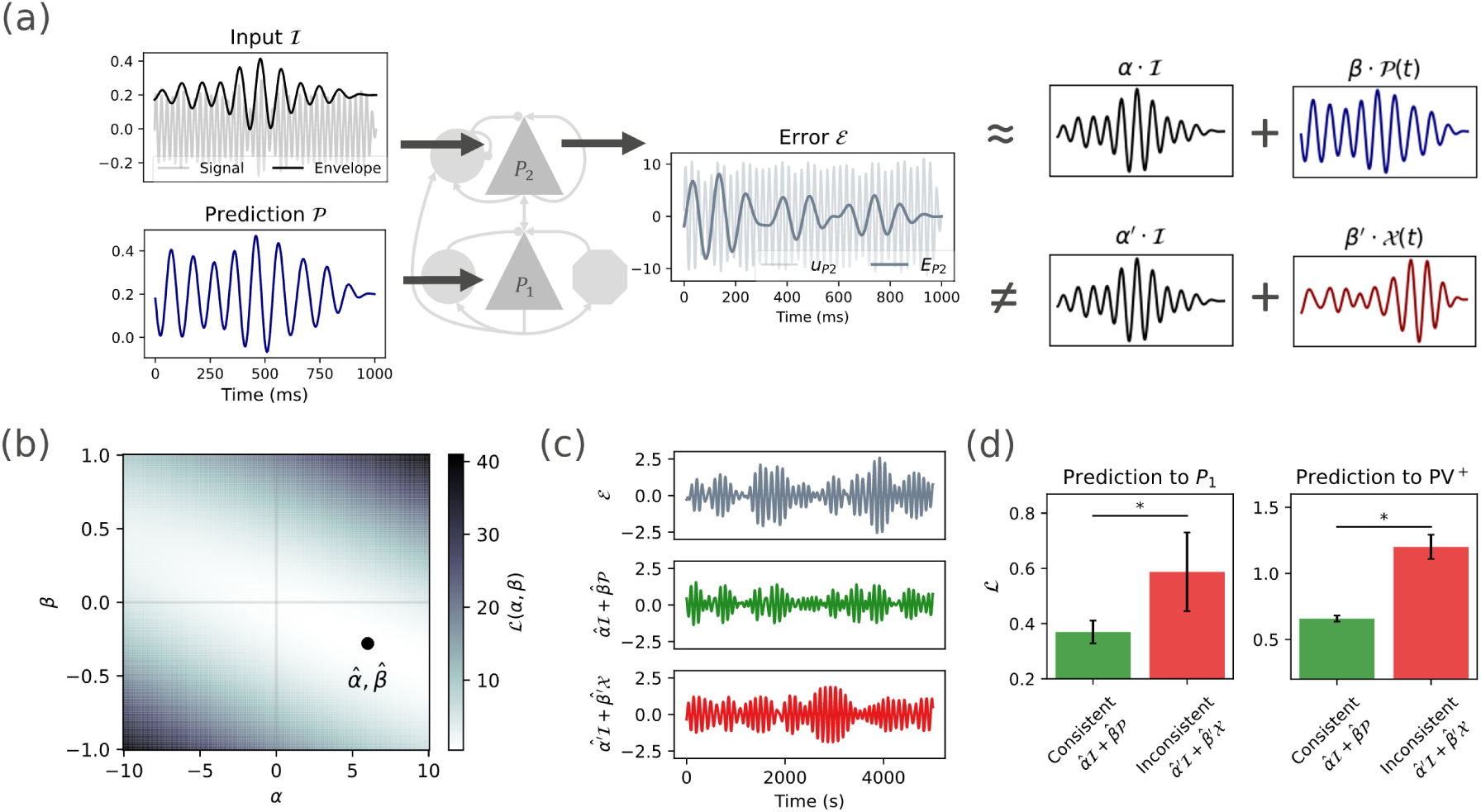
SEC subtraction scheme, prediction sent to *P* 1. (a) We can find a linear combination of the input and prediction signals *α* · I(*t*) + *β* · P(*t*) that approximates the error signal, with *α* and *β* of different sign (subtraction). Using an unrelated signal X(*t*) for the linear combination should lead to a worse approximation. (b) Optimal values *α*^*, β*^^^ for a specific pair (I, P). The loss function L is defined as the MSE between the error signal and the linear combination of input and prediction. (c) The error computed by the LaNMM (grey traces) is well approximated by the optimal subtraction of input and prediction (green traces) but not by the optimal subtraction of the input and an unrelated signal (red traces). (d) The optimal subtraction using I and P (consistent case) is closer to the LaNMM output than the optimal subtraction using an *I* and an unrelated signal X (inconsistent case). Prediction signal sent to *P*_1_ (left) or PV (right).

### 3.3 EEC subserves Slow-Time Precision Modulation

Figures 6 depicts how EEC suppresses the precision of error signals: the error signal is significantly attenuated (lower error power) when using a precision-modulating signal with transient increases in alpha power (slow envelope), as opposed to the case where a flat, non-modulating prior is sent to *P* 1.

Figure 8 illustrates how EEC in the LaNMM can be used to perform modulation of prediction errors by subtraction. The error signal precision encodes the difference between the input precision and the precision-modulating prior signal, as quantified by the MSE between the normalized error precision and the subtraction of the Π_I_ and Π_P_. The similarity between the LaNMM output (E precision, Π_E_ ) and the subtraction of input and prior precisions (Π_I_ − Π_P_) is illustrated in Figure 8b. Overall, the results indicate that EEC can be harnessed to modulate the precision of error signals.

**Figure 8:**
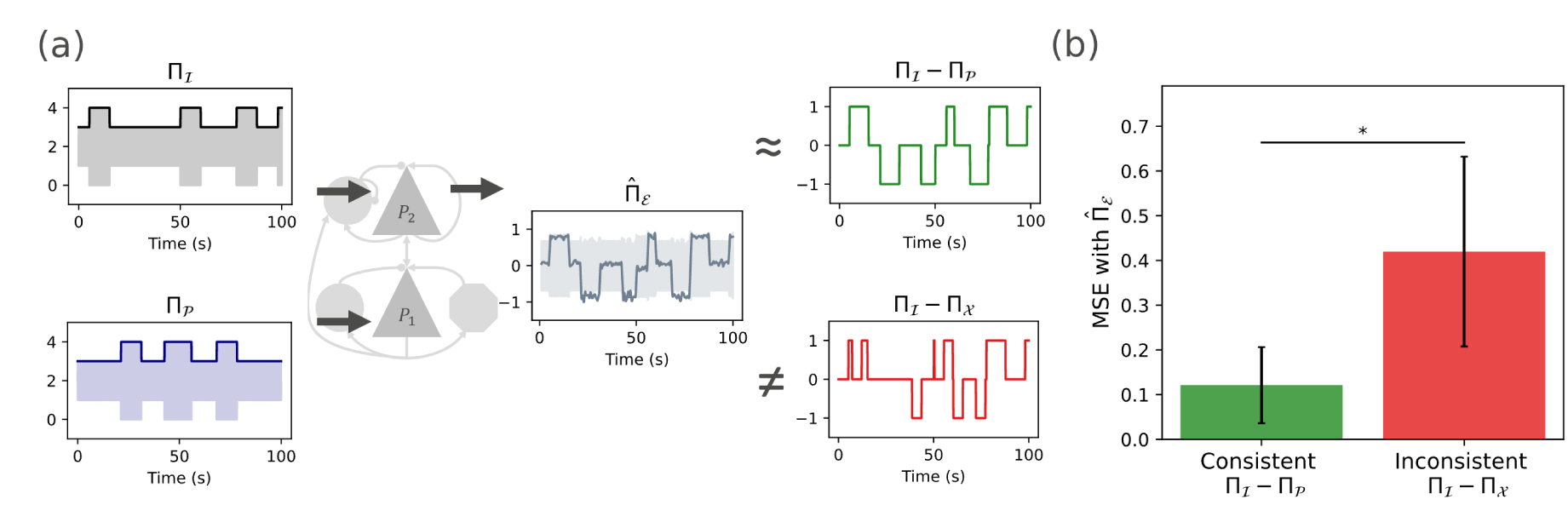
EEC subtraction scheme. (a) The error signal precision encodes the difference between input precision and prior precision. (b) For a given input and prior encoding some precision modulation sent to the LaNMM, the output error precision is better approximated by the difference Π_I_ − Π_Π_ (consistent case) than by the difference between the input precision and unrelated modulation-precision signal Π_I_ − Π_X_ (inconsistent case), as captured by the MSE between the LaNMM output Π^^^_E_ and the subtraction of each case.

### 3.4 Psychedelics and Interneuron Pathology Disrupt the Comparator

Finally, we investigated how LaNMM disruptions associated with serotonergic psychedelics or Alzheimer’s Disease interneuron pathology impact the comparator functions.

#### 3.4.1 Serotonergic Psychedelics

Figure 9 illustrates how the increased excitability due to serotonergic psychedelics, reflected in the model by an increase of the parameter *A_P_*_1_ , translates into power spectrum alterations and the disruption of the comparison process. In particular, we can see how the low prediction error resulting from a matching input and prior in the baseline condition (low *P*_2_ envelope power) is increased in the psychedelics condition (Figures 9c and 9e), resulting in an error similar to the one obtained when a flat prior is applied (see Figure 5a). Moreover, under the psychedelics condition, the influence of the precision attenuation signal is greatly reduced, causing the error to increase and primarily reflect the input signal (Figures 9d and 9f).

**Figure 9:**
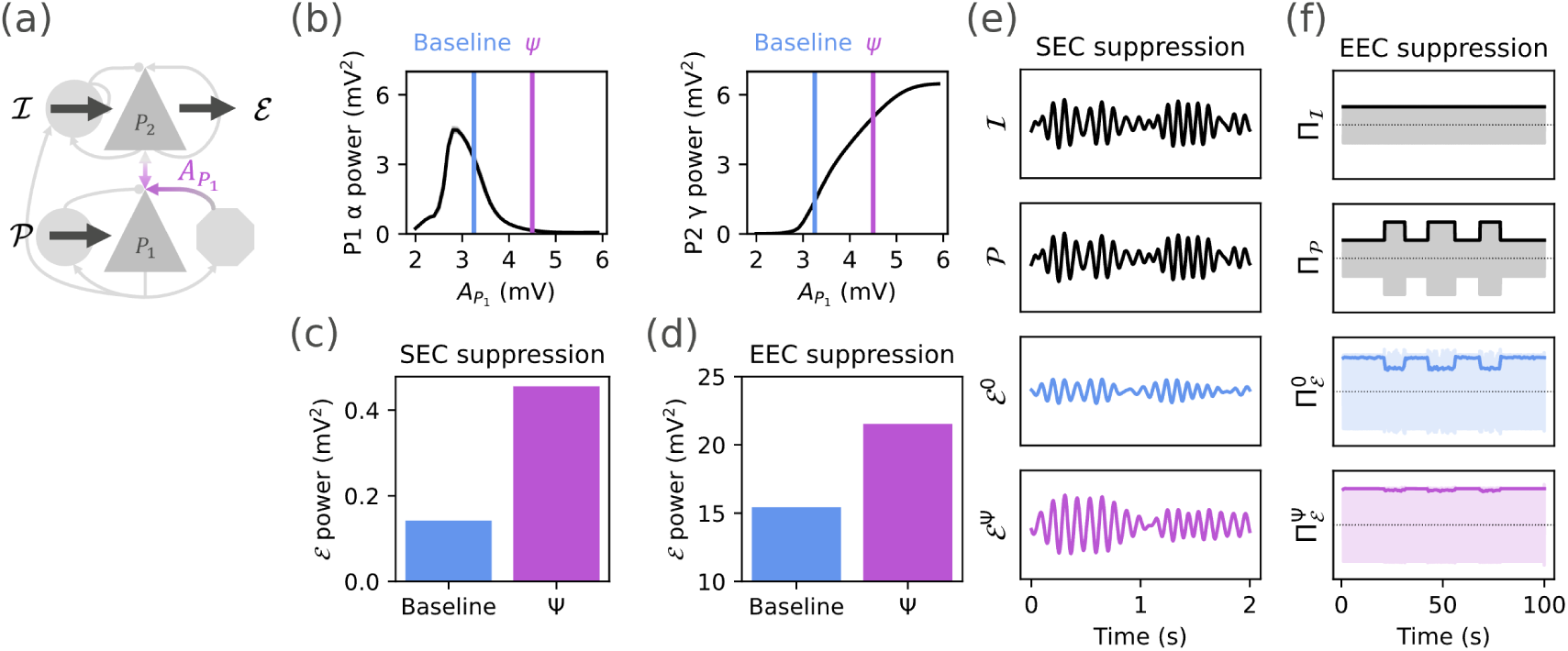
Effects of serotonergic psychedelic alterations on the LaNMM model. (a) The excitatory effects of serotonergic psychedelics are implemented as an increase of the excitatory synaptic gains of the *P*_1_ population, *A_P_*_1_ .^97^ (b) The alteration of *A_P_*_1_ in the model leads to decreased alpha power and increased gamma power. (c) For the SEC mechanism, the error suppression for a prediction matching the input is altered by psychedelics, resulting in a higher power of the error signal. (d) For the EEC mechanism, the influence of the precision-modulating signal is diminished, also resulting in increased error. (e, f) Example signals illustrating the SEC and EEC comparator mechanism for the baseline and psychedelics condition, showing a reduced effect of the prior and precision-weighting modulation in the psychedelics condition.

Overall, these results suggest that serotonergic psychedelics increase the effective weight of inputs due to increased power of fast (gamma) oscillations, and diminish the effect of predictions and precision-modulating signals, as a consequence of the reduced power in slow (alpha) oscillations. This results in higher prediction error signals in the psychedelics condition.

#### 3.4.2 Alzheimer’s Disease

Figure 10 illustrates how positive SEC/negative EEC patterns and comparator functionality degrade under AD-disrupted predictive coding conditions. The alteration of the *C_P V_* _→_*_P_* _2_ parameter leads to an increase in alpha and gamma power in early stages (hyper-excitability) and an overall power decrease in late stages.

**Figure 10:**
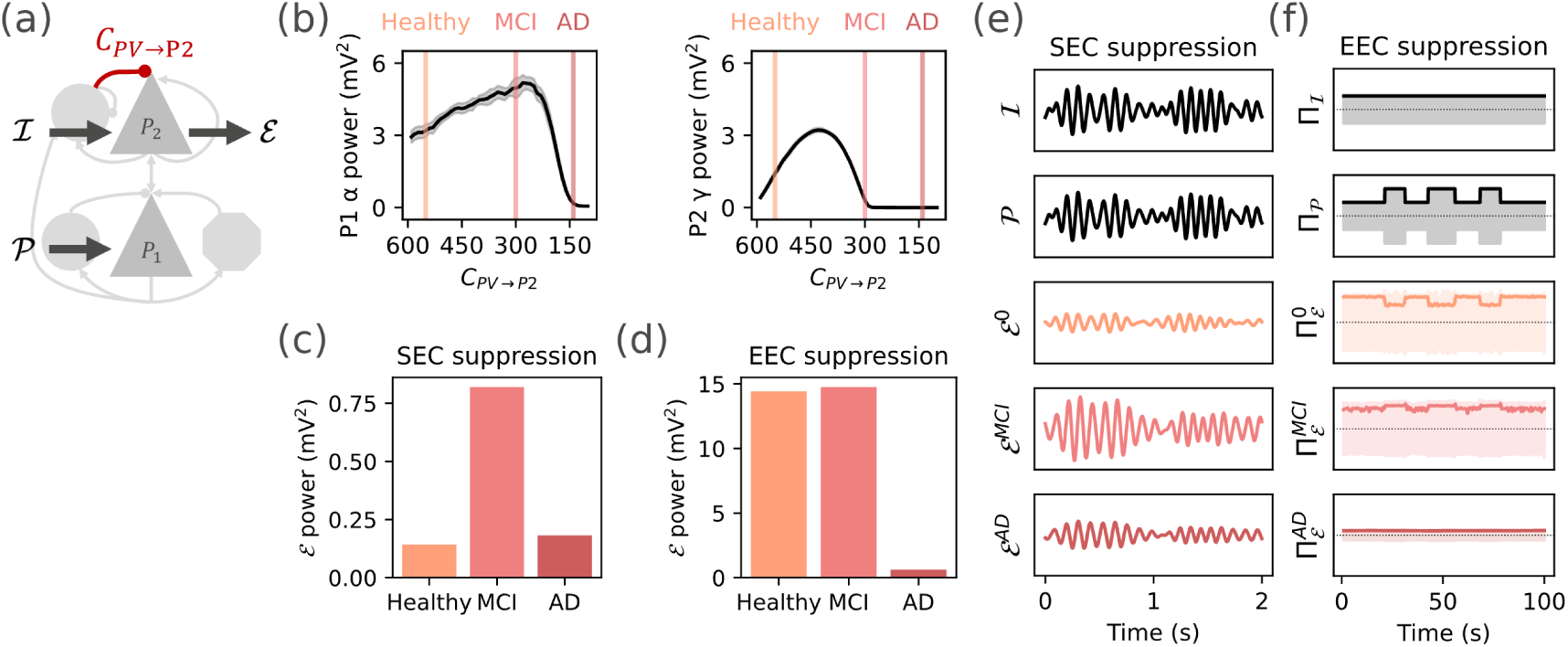
Effects of AD-like pathological alterations on the LaNMM model. (a) AD pathology is introduced by reducing the connectivity of the PV to *P*_2_ population, *C_P V_* _→_*_P_* _2_.^79^ (b) The alteration of *C_P V_* _→_*_P_* _2_ reflects the different stages in the progression to AD and leads to an increase in alpha and gamma power in early stages and an overall power decrease in late stages. (c) The suppression of prediction error observed in the healthy stage for matching input and prediction is disrupted in AD stages (especially MCI), resulting in higher prediction errors. This can be explained by a reversal of sign in SEC (see Figure H1). (d) The effect of the precision-modulating signal is reduced in the MCI stage, resulting in higher error power, while in the AD stage, the error power is substantially diminished. (e, f) Sample signals illustrating the SEC and EEC comparator mechanism for the baseline and altered conditions, showing a reduced effect of the prior and precision-weighting modulation in the altered condition.

In both the MCI and AD stages, we find that the error evaluation process is disrupted with respect to the healthy stage. In particular, the low prediction error obtained through SEC for a matching input and prior is strongly increased in MCI stage and then slightly increased in the AD stage with respect to the healthy condition (Figure 10c). The low prediction error power observed in AD reflects Comparator dysfunction: the similarity in error power between AD and healthy conditions arises not from accurate prediction, but rather from an overall reduction in cortical power typical of AD, as illustrated by decreased alpha and gamma activity in Figure 10b.

This is confirmed by the case of the EEC mechanism—panels (d) and (f)—, where the effect of a precision-modulating signal is reduced in the MCI condition (higher prediction error as well), but the error power is drastically reduced in the AD stage (Figure 10f).

In sum, these results suggest that in the early stages of AD, error evaluation and precision are disrupted (increased prediction error computation and reduced influence of precision-modulating signals), leading to higher prediction errors even when signals match. In later stages, prediction errors are abnormally suppressed regardless of predictions or the modulation of precision by top-down signals.

## 4 Discussion

### 4.1 Interpretation of results

Our modeling results show that it is possible to implement the evaluation of E in various ways using the cross-frequency properties of the LaNMM, with an information encoding scheme where information is stored in signals or in their envelopes.

We first showed how the model, both in its nominal parametrization^105^ and in a wider mean input landscape, provides robust bidirectional cross-frequency coupling: controlling the input fluctuations into *P*_1_, *P*_2_ or PV results in envelope fluctuations in the other population (*P*_2_, *P*_1_, and PV, respectively) (see Figure 4). More specifically, we observed robust positive SEC, consistent with the previously reported PAC.^105, 129^ With regard to negative EEC, it is noteworthy how the two subpopulations operate in what we may call *dynamical competition*, a sort of zero-sum game of the total power generated by the column. Cross-frequency coupling provides the basis for the rest of our results, and, in particular, SEC motivates the further exploration of an AM information encoding scheme in neural signals.^120, 130^

Exploiting SEC and EEC, we then showed how the evaluation of E can be carried out by the laminar column. In the first case, information carried by the fast oscillation envelope arriving at *P*_2_ is contrasted with a slow prediction signal, producing effective suppression if the prediction is accurate. Two mechanisms were provided. The first was implemented with the signal predictions arriving at *P*_1_ with a phase shift of around 180° with respect to the input signal. In the second implementation, predictions arrived at the PV cell in the PING subpopulation in phase with the input signal.

In the case of EEC, where information about precision is carried by the “envelope of the envelope” of the fast oscillation at a slower time scale, the prior precision is encoded as the envelope of the slower signal arriving at *P*_1_, without phase shift or sign reversal, and the precision modulation is carried out over a longer time scale (determined by the time scale of the envelope of the slow signal). Figure 3 illustrates this scenario.

An instructive analogy with regard to the role of time scales comes from radar signal processing, particularly Synthetic Aperture Radar (SAR), which distinguishes between *fast-time* and *slow-time scales*.^131^ In SAR, fast-time corresponds to the rapid reflections of individual radar pulses used for high-resolution range measurements—analogous to rapid signal-to-envelope coupling (SEC) for fast prediction-error computations. Conversely, slow-time relates to the slower progression of the radar platform across multiple pulses, enabling integration of spatial information over a larger scale—mirroring envelope-to-envelope coupling (EEC), which facilitates predictive integration and gating at slower timescales.

### 4.2 Predictive coding and precision

Our results with EEC provide a mechanism for neural processing proposals hinging on the availability of gating, which is necessary for the Bayesian implementation of predictive coding. Technically, predictive coding was introduced in the 1950s as a scheme for compressing sound files.^132^ This is conceptually pertinent to KT in the sense that compression is exactly the objective prescribed by minimizing algorithmic complexity under this framework. In the neurosciences, predictive coding was subsequently applied to continuous (neuronal) signals and state estimation in the brain.^1, 20, 133^ From an engineering perspective, predictive coding for continuous state space models corresponds to Bayesian filtering and, in the limit of linear models, Kalman filtering.^1^ The Kalman filtering perspective is important because it reveals that there are two things that need to be estimated in predictive coding: the prediction error *per se* and the (Kalman) gain that is to be applied to the prediction error for model update (see Appendix D). Crucially, the Kalman gain itself has to be optimized, which requires an estimation of the precision (i.e., inverse variance) of sensory and state noise. In short, there are two kinds of elements that need to be predicted in predictive coding: the (first-order) content and the (second-order) precision. This dual aspect inherits directly from the assumption of Gaussian posterior estimates in predictive coding, which have two sufficient statistics, namely, (first-order) mean and (second-order) precision (inverse covariance).

When translating this dual estimation problem into the frequency domain, the current formulation associates this with cross-frequency coupling of a fast and slow sort, respectively (i.e., phase-amplitude or, more generally, SEC, and amplitude-amplitude or EEC coupling). Intuitively, assigning a high precision to a prediction error enables it to have a greater effect on subsequent state estimation or (Bayesian) belief updating. This manifests—in deep or hierarchical models—as an optimal gating of ascending prediction errors, such that precise prediction errors are selected over imprecise prediction errors. When the precision of prediction errors fluctuates, then the implicit selection, routing, or gating acquires a dynamic aspect, resulting in a Bayes-optimal dynamic coordination, routing or gating.

### 4.3 Gating by Inhibition and Predictive Routing

As we have seen, EEC can support *gating*. Gating mechanisms are used to implement fundamental logic gates, the building blocks of all digital computation and self-regulating systems.^134^ Gates lie at the heart of every computational system because they enable conditional branching—the capacity to direct information flow according to context. While Leibniz’s Step Reckoner^135^ and Babbage’s Difference Engines relied on mechanical gears and levers as switches for arithmetic operations,^136^ the introduction of vacuum tubes in Colossus enabled high-speed gating for codebreaking.^137^ Today, modern computers integrate billions of transistor-based switches for reprogrammable logic.^138^ By contrast, the brain accomplishes switching via inhibitory gating, where interneurons toggle neural pathways by hyperpolarizing downstream targets—or, as discussed here, through competitive oscillations via EEC.

In the context of predictive coding, although we have focused on the columnar implementation of the Comparator with information encoded in temporal signals, the brain also encodes information in space. That is, not only *when* but also *where* the signals arrive in the network matters. In the case of local error evaluation, weighted error predictions must arrive at the right location to be effective in suppressing the propagation of errors. That is, agent predictions are in general of the form “these signals will arrive at this location at these times”. This expectation may be an active one, where the system reconfigures (via switches, for example) to route information streams in a particular way.

This gating provides the same conditional-branching capability seen in electronic hardware: specific signals are allowed or blocked depending on context, supporting attention, learning, and adaptive decision-making.^139^ Thus, whether realized with mechanical gears, vacuum tubes, transistors, synaptic inhibition, or oscillatory competition, switching at different spatiotemporal scales is foundational to any system that must selectively route information—and it underscores the shared computational logic between engineered machines and biological neural networks.

In contrast with the local evaluation of prediction error, *routing*^22^ *or gating*^21^ refer to the dynamic modulation of information flow across cortical hierarchies, selectively directing or suppressing signal propagation. Gating mechanisms determine which signals can pass through at any given time and modulate the gain of signals based on relevance or salience. This is sometimes referred to as *gating by inhibition*^21^, a framework where optimal task performance correlates with alpha activity in task-irrelevant areas. Gating by inhibition has been proposed as an attentional mechanism, where the agent chooses what information streams to process by selectively dampening or enhancing signals through inhibitory processes.^21^ In modern neural network transformer architectures, a similar principle of selective attention is implemented via multi-head attention, which learns to weight different input elements, effectively gating the flow of information in deep networks.^140^

*Predictive routing* is gating-related proposed mechanism where top-down predictions dynamically shape sensory information flow by selectively inhibiting expected inputs through amplitude-amplitude coupling (AAC),^22^ the equivalent of EEC in our framework. In predictive routing, the brain anticipates incoming sensory data based on prior knowledge and actively suppresses the cortical pathways associated with these predictions. This can be interpreted in light of prediction error evaluation, where selective inhibition implements the Comparator in a distributed manner (not on a single column) and on a longer, coarse-grained timescale than SEC, dynamically routing unexpected signals for further processing. In this context, *information is encoded by route* and predictions address the route taken by sensory stimuli: correctly predicted inputs are inhibited, while unpredicted inputs propagate, signaling prediction errors (see Appendix E and Figure E1 for further details). Top-down predictions are communicated via alpha/beta oscillations (8–30 Hz) originating from deep cortical layers. These rhythms prepare sensory pathways by pre-emptively inhibiting the columns that process expected inputs. It is this preemptive aspect that reflects predictions of which prediction errors will convey precise or salient information.

The interplay between these dynamics, governed by the laminar organization of the cortex, ensures that deep layers provide the inhibitory feedback while superficial layers relay the feedforward sensory information. This layer-specific interaction dynamically selects the route for information flow. Thus, EEC provides a mechanism for weighted spatiotemporal information processing of prediction errors as well as other information processing agent choices in network space and time (e.g., attention and sensory attenuation). In all these instances, it reflects the choice and optimal use of *relevant information* for model updates conducive to the maximization of the agent’s Objective function.

### 4.4 Error Evaluation Disruption

Our results provide a biologically grounded model for CFC and error evaluation in the cortical column and a measure of their disruption under the effects of psychedelics or PV alterations in the brain. Here, we connect our modeling results to experimental findings on CFC and prediction error evaluation in these conditions.

#### 4.4.1 Serotonergic Psychedelics

The present work expands on our previous research on the effect of serotonergic psychedelics.^97^ Our results indicate that serotonergic psychedelics weaken the influence of predictions and precision attenuation mechanisms on error computation, leading to increased (precision-weighted) error signals even when the prediction matches the input. The reduced influence of top-down precision-modulation signals suggests that, under psychedelics, the error signal primarily reflects the input itself rather than the mismatch between input and prediction, effectively suppressing the prediction’s influence (Figure 9). This computational shift could explain psychedelic-induced perceptual distortions, where sensory signals are experienced with reduced interpretative constraints, as proposed by the REBUS model.^91^ By disrupting hierarchical inference in this way, psychedelics may both destabilize perception and facilitate the revision of rigid priors, providing insight into their potential therapeutic effects.

In empirical studies, the relaxation of top-down constraints under psychedelics manifests in oscillatory changes in MEG/EEG, notably a broadband desynchronization of cortical rhythms.^92^ This global desynchronization is thought to index an increase in neuronal entropy or disorder^91^—essentially, neurons firing in a less constrained manner. In terms of predictive coding, such desynchronization corresponds to a collapse of precise priors (organized oscillatory states) and a release of sensory processing from descending constraints. The alteration of gamma power and the reduction of alpha oscillations under psychedelics^141, 142^ likely contributes to this: since cross-frequency coupling often involves alpha/beta phase gating gamma, the disruption of slower oscillations means gamma activity is no longer properly coordinated in time. Consequently, it has been suggested that the alpha-gamma and theta-gamma coupling that orchestrates hierarchical processing is weakened under psychedelics, and with it, the influence of sensory prediction errors increases.

In our model, the changes in excitability due to serotonergic psychedelics lead to oscillatory changes in the cortical column (Figure 9b) that increase the effective weight of inputs (increased gamma power) and diminish the *effect* of arriving predictions (decreased alpha power), resulting in higher prediction error signals. Our results further suggest that the increase of prediction error signals traveling to upstream hierarchical layers may translate into a *reduction of precision* of top-down priors from upper layers (e.g., in the alpha band). Although our framework does not address how prediction error signals are translated into model and model precision updates, the above is consistent with a mathematical implementation of model updates (e.g., Kalman filtering), where the uncertainty in incoming data impacts confidence in the model.

#### 4.4.2 PV Interneuron Dysfunction in Alzheimer’s, Schizophrenia, and ASD

We extended our earlier LaNMM model of AD^79^ by connecting the pathology in AD with information processing dysfunction, demonstrating how predictive coding mechanisms are progressively impaired as the disease advances. Our results indicate that these progressive alterations in PV interneuron function lead to disruptions in both SEC and EEC mechanisms, affecting the brain’s ability to suppress and compute prediction errors.

In the preclinical phase of AD—prior to mild cognitive impairment (MCI)—individuals often report Subjective Cognitive Decline (SCD), defined as self-perceived worsening of cognitive function despite normal performance on standard tests.^143^ SCD is frequently one of the earliest detectable changes in the AD continuum, preceding measurable deficits and dementia. Consistent with our model, SCD may reflect an early increased flow of prediction errors. Indeed, in our simulations of the MCI stage, the increased alpha and gamma power contribute to heightened error signals, reflecting an overactive yet unstable predictive system. The disruption of error computation is further highlighted by the reduced influence of precision-modulating signals (Figure 10f), revealing a severe breakdown of gating mechanisms in both MCI and AD stages. Notably, the largest disruptions are observed in MCI, consistent with the idea that interneuron dysfunction and alpha/gamma hyperactivity initially drive pathological changes before leading to widespread network collapse.

In the clinical phase of AD, the auditory MMN is significantly reduced compared to healthy aging.^70^ This attenuation of MMN is especially pronounced at longer inter-stimulus intervals, suggesting a rapid decay of the “memory trace” for standard stimuli—i.e. an impaired ability to maintain predictions over time. Even individuals at risk for AD (e.g., APOE4 carriers) show smaller MMNs than controls before clinical symptoms emerge.^70^ AD also impairs the detection of higher-order regularities; for example, patients show deficits in perceiving melodic contour violations (sequence patterns in music),^70^ indicating trouble with global predictive inferences. At a later processing stage, the P300 component is delayed and diminished in AD. Numerous studies and meta-analyses have found prolonged P300 latency and reduced amplitude in AD patients, correlating with cognitive decline.^71, 72^ These ERP biomarkers suggest that AD brains generate weaker prediction errors and/or fail to effectively propagate them to higher levels for conscious detection. In our model, in the AD stage, the overall power decrease—particularly in high-frequency bands—reduces the ability of the system to generate reliable prediction errors, leading to a collapse of error-signaling and comparator functionality. This is evident in the SEC scenario, where error suppression appears to recover in AD but is actually a byproduct of overall power loss rather than accurate predictive coding (Figure 10c).

Finally, fast-spiking PV interneurons, essential for inhibitory control and gamma synchronization, are also disrupted in other neuropsychiatric conditions, including ASD,^83^ ADHD,^85, 86^ and schizophrenia.^84^ Our model of PV dysfunction introduced in Sanchez-Todo (2025),^79^ and extended here in the context of predictive coding, is therefore not only applicable to AD but may also capture core mechanisms underlying these disorders, all of which exhibit deficits in information processing, including altered sensory attenuation.^80–82, 88–90^

### 4.5 Previous Computational Models of CFC and Prediction Error

Several physiologically grounded neural mass and network models have been developed to reproduce CFC phenomena such as PAC and AAC. For instance, Mejias et al. (2016)^127^ introduced a laminar cortical network model of the primate visual hierarchy that captures distinct frequency-specific feedforward vs. feedback signaling. Slow oscillatory input from deep layers modulated superficial gamma, yielding PAC between layer-5/6 alpha (∼10 Hz) and layer-2/3 gamma (30–70 Hz). At the mesoscopic scale, NMMs with realistic neuronal interactions can intrinsically generate PAC. Chehelcheraghi et al. (2016)^110^ adapted a cortical column model with pyramidal cells and fast and slow inhibitory interneurons. Under noise, the fast inhibitory subpopulation became a self-oscillatory gamma source, while the slow inhibitory rhythm in the alpha–theta range modulated the gamma amplitude, producing robust PAC. Extensions of this model also demonstrated other coupling types (phase–frequency, amplitude–amplitude, frequency–frequency). Sotero (2016)^144^ highlighted how connectivity structure in a cortical column NMM shapes PAC generation, with balanced excitation–inhibition and resonant synapses modulating cross-frequency interactions. More recently, “next-generation” NMMs^145, 146^ replicated canonical CFC phenomena in coupled oscillatory circuits, attributing PAC emergence to interacting inhibitory populations or slow sinusoidal drives near a Hopf bifurcation. At the spiking-neuron level, models have also captured CFC with explicit physiological mappings.^147^ Meanwhile, thalamocortical models showed how cortical slow oscillations (∼1 Hz) modulate thalamic sleep spindles (12–15 Hz), again reflecting phase-amplitude relationships.^148^

In parallel, some computational models explicitly encode prediction errors within physiologically grounded architectures. Rao and Ballard^1^ proposed a hierarchical visual model where higher-level feedback suppresses predicted inputs, forwarding only the residual unexplained signal. Spratling’s cortical models^149^ similarly use inhibitory feedback for residual error computation, and a network by Wacongne et al.^150^ reproduces mismatch negativity by comparing predicted vs. actual tones. Canonical microcircuit models further propose that distinct cortical populations encode predictions vs. errors (deep vs. superficial layers), though details of how the error is evaluated remain less explicit.^3^ Song et al.^151^ extended these ideas to a pain perception NMM in which top–down analgesic expectations modulate incoming nociceptive signals, generating a residual error that drives cortical responses.

Though these studies illustrate how either CFC or explicit prediction error signals can be implemented, most do not address how cross-frequency interactions themselves underwrite prediction errors. In contrast, our approach incorporates both the generation of CFC and its computational role. By leveraging SEC and EEC, the proposed LaNMM provides a physiologically grounded instantiation of the Comparator function central to predictive coding frameworks. Unlike prior models that rely on noise-driven dynamics or impose CFC via structural constraints, the LaNMM produces emergent error signals approximating the difference between bottom-up sensory inputs (fast oscillatory envelopes) and top-down predictions (slower oscillations). This biologically plausible mechanism enables the dynamic, hierarchical modulation of prediction errors across multiple timescales. Furthermore, by integrating physiological constraints on laminar-specific processing, the LaNMM offers a unifying framework for studying predictive coding across cortical layers. In doing so, it builds a mechanistic bridge between abstract predictive coding theories and the neurophysiological generation of cross-frequency couplings, yielding new insights into both normal and pathological brain function.

### 4.6 Limitations

While our LaNMM approach captures key features of cross-frequency coupling and error evaluation, several limitations must be acknowledged. First, the model is an abstraction that simplifies complex cellular and synaptic dynamics; parameters were chosen based on theoretical and empirical considerations but may not fully capture the heterogeneous properties of real cortical networks.^3, 4^ Second, although our simulations reproduce oscillatory signatures observed in Alzheimer’s disease and psychedelic states, such as altered gamma power and CFC patterns,^91, 152^ the direct translation of these findings to *in vivo* conditions requires further validation through combined EEG/MEG and invasive recordings. Third, the impact of neuromodulatory systems—especially the cholinergic and serotonergic influences—has been modeled in a simplified manner, and future work should integrate more detailed receptor-level dynamics to better account for state-dependent changes in error signaling. Fourth, our analysis covers only mesoscopic processing in single-node dynamics, while both micro and macro network effects are undoubtedly relevant in predictive processing.

Finally, we have focused mainly here on the analysis of prediction error evaluation at the single column level and using a particular model displaying oscillations in the alpha and gamma bands. The model could be adjusted to other frequencies through parameter changes, and indeed, other models of oscillations exhibiting CFC could be used for the same purpose. Moreover, the framework proposed can be extended across spatial and temporal scales. In particular, the information encoding scheme based on amplitude modulation can be extended hierarchically, with information encoded in signals, their envelopes, the envelopes of envelopes,^120^ etc.

## 5 Conclusion

Our work supports the view that cross-frequency coupling provides a functional substrate for the evaluation of weighted prediction errors within a hierarchical predictive coding framework. The model presented here demonstrates how distinct modes of amplitude modulation—signal-envelope and envelope-envelope coupling—can instantiate a Comparator mechanism leveraging fast (gamma) and slow (alpha/beta) oscillatory dynamics. Moreover, the model speaks to causal relationships between neural CFC and cognitive functions as well as a mechanistic perspective on how pathophysiology, such as in AD or induced by serotonergic psychedelics, disrupts the balance between top-down predictions and bottom-up error signaling. Further work may explore the implications in other conditions associated with predictive coding abnormalities, such as schizophrenia, ASD, or ADHD. Our mechanistic insights not only deepen our understanding of multiscale information processing in the brain or other systems relying on oscillatory computation but can also guide the development of potential therapeutic interventions aimed at restoring normal oscillatory function and predictive coding.

## Funding

Giulio Ruffini, Edmundo Lopez-Sola, Francesca Castaldo, Roser Sanchez-Todo, are funded by the European Commission under European Union’s Horizon 2020 research and innovation programme Grant Number 101017716 (Neurotwin) and European Research Council (ERC Synergy Galvani) under the European Union’s Horizon 2020 research and innovation program Grant Number 855109. Jakub Vohryzek is funded by Neurotwin (855109).

## Conflict of Interest

GR, ELS, RST and FC work for Neuroelectrics, a company developing computational brain stimulation solutions for neuropsychiatric disorders.

## A Band power and PEIX (saturation) plots

Figures A1 and A2 illustrate how power distributions within each frequency band are influenced by the mean external inputs to the respective populations. We also display the Population Excitation-Inhibition Index (PEIX), a normalized slope measure of excitation–inhibition balance.

## A.1 PEIX computation

Neural populations are often modeled by a sigmoid function,

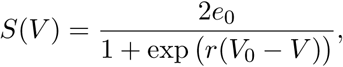

where V0 is the half-maximum potential (the linear operating point), e0 is the maximum firing rate, and r sets the slope. The derivative,

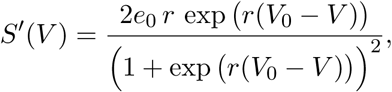

peaks at V = V0 with 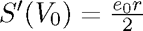 and decreases as the system enters its excitatory (V ≫ V0) or inhibitory (V ≪ V0) saturation regimes.

To quantify the deviation from the ideal linear regime, we define the dimensionless variable

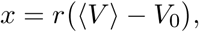

where ⟨V ⟩ is the average membrane potential. By normalizing the local slope S′(V ) with its maximum value at V0, we obtain a measure of nonlinearity that distinguishes between excitation and inhibition. The PEIX is defined as

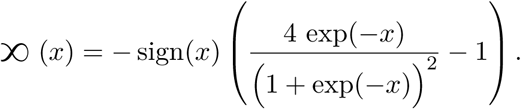

This formulation ensures that:

• (x) ≈ 0 when ⟨V ⟩ ≈ V0 (linear regime),

• (x) > 0 when ⟨V ⟩ > V0 (excitatory state),

• (x) < 0 when ⟨V ⟩ < V0 (inhibitory state),

• | (x)| ≈ 1 in the saturated regimes.

Thus, by focusing on the normalized slope of the sigmoid, PEIX provides a compact, dimensionless index of both the deviation from the optimal operating point and the underlying excitation–inhibition balance.

## B Gain optimization

In this section, we detail the methodology used to select the gains applied to the input and prior messages provided to the LaNMM for the SEC and EEC error computation.

In each case (SEC / EEC), we first selected the input gain GI so that when the input message was applied alone (no prediction signal), the output of the LaNMM (E) showed a high PCC with the input message. This way, we could ensure that the input was strong enough to modulate the activity of the LaNMM and carry it through the output message. As shown in Figure B1, if the input gain is sufficiently high, the output is highly correlated with the input. However, the higher the input gain, the more it would dominate the output message and the harder it would be to suppress it with a prior message. We thus selected in each case the lowest input gain that resulted in high PCC with the input, i.e., G∗ (SEC) = 10 and G∗ (EEC) = 1.

For the SEC mechanism, we used the selected input gain and evaluated the E power as a function of the prior gain in the case when a prior message matching the input was sent to the LaNMM. We studied the cases where the prior was sent to the P1 population and to the PV population. In each case, there was an optimal value of prior gain that would lead to a minimum error, as shown in Figure B1. We chose the nearest round values that better approximated such a minimum, i.e., G∗ (SEC, P 1) = 20, G∗ (SEC, PV ) = 5.

For the EEC mechanism, we used the selected input gain and evaluated for different prior gains the MSE between the subtraction of input precision and prior precision and the error precision, as shown in Figure 8. We selected the nearest round value of prior gain that resulted in the minimum MSE, G∗ (EEC) = 10, as shown in Figure B1.

## C Input-prior phase dependence (SEC)

We expected that for effective suppression of the input signal with a matching prior using the SEC mechanism, the prior signal sent to P1 had to be phase-shifted by 180° (i.e., inverted). This inversion would allow the prediction message to counteract the excitatory input effectively. This was due to the fact that P1 is an excitatory population and that the SEC for P1-driving inputs is positive, as shown in Figure 4. We confirmed this by computing the power of the output from the LaNMM E when varying the phase of the matching prior (see Figure C1a). Interestingly, we can see that when applying the matching prior in phase with the input (0° shift), the output error is increased with respect to the flat prior and wrong prior conditions.

In contrast, for the PV population, which is inhibitory and has positive SEC for PV-driving inputs, the prior had to be applied in phase with the input signal to achieve suppression (see Figure C1b. When priors shifted 180° (inverted) were sent to the PV population, the LaNMM output E power was increased.

## D Precision and Kalman Gain

In the Kalman filtering framework, updating the state estimate involves balancing prior predictions with incoming measurements. At step k, this balance is quantified by the innovation covariance Sk, defined as:

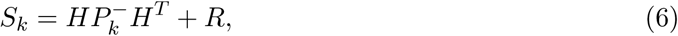

where:

• P − is the predicted (prior) state covariance, reflecting the uncertainty inherent in the internal model’s prediction of the current state before incorporating new observations.

• H is the observation model, projecting uncertainty from state-space into measurement-space, transforming prior state uncertainty P − into observation uncertainty HP −HT .

• R is the measurement noise covariance, explicitly encoding uncertainty or noise inherent to the measurements themselves, independent of the model.

Thus, the innovation covariance Sk combines two distinct sources of uncertainty:

1. Model-based uncertainty (HP −HT ): uncertainty arising from the prediction step, reflecting imprecision in the model’s prior expectation about the observed quantities.

2. Measurement uncertainty (R): intrinsic uncertainty in the data, reflecting how noisy or unreliable the incoming observations are.

The precision matrix Πk, defined as the inverse of Sk, explicitly integrates both these uncertainties:

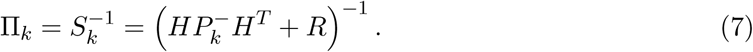

Higher precision thus means lower combined uncertainty, increasing confidence in the prediction errors.

The Kalman gain Kk, directly related to precision, weights prediction errors (innovations) when updating the state estimate:

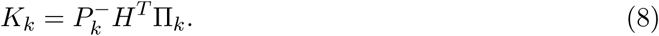

Finally, the posterior state estimate x^+ is updated using this gain applied to the prediction error νk = zk − Hx^−:

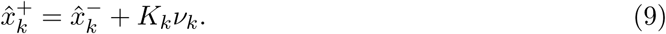

Precision thus clearly separates and balances model-derived and measurement-derived uncertainties, directly controlling the integration of observational evidence into state inference.

## E Predictive Routing from EEC

In predictive routing,22 information is encoded by route and predictions address the route taken by sensory stimuli: correctly predicted inputs are inhibited, while unpredicted inputs propagate, signaling prediction errors.

Top-down predictions are communicated via alpha/beta oscillations (8–30 Hz) originating from deep cortical layers. These rhythms prepare sensory pathways by pre-emptively inhibiting the columns that process expected inputs. It is this pre-emptive aspect that reflects predictions of which prediction errors will convey precise or salient information.

When a prediction specifies a particular route—say, Route A—the corresponding cortical column is suppressed by alpha/beta activity, which in turn reduces gamma oscillations and neuronal spiking in that column. As a consequence, expected stimuli arriving via Route A are effectively blocked. Conversely, when the prediction is off (inhibition of Route B), the stimulus bypasses the inhibitory control because the predicted pathway is not engaged, generating an active output. In this case, the absence of alpha/beta-mediated inhibition allows for enhanced gamma oscillations and increased spiking activity, which serve as a robust signal of a prediction error. Figure E1 illustrates this process: when predictions indicate that a stimulus will follow Route A, alpha/beta rhythms suppress gamma activity in the corresponding cortical column, blocking that pathway. If the actual input follows Route B, the lack of inhibition permits gamma activity and spiking, thereby signaling a prediction error and routing unpredicted information upward in the cortical hierarchy. This provides a distributed implementation of the Comparator using multiple columns.

## F LaNMM examples in other bands

Here we provide a couple of examples of the spectra produced by other parameter configurations in LaNMM, obtained by changing synaptic time constants.

## G Input generation for SEC and EEC characterization

Generation of Multiscale Noise Signals. The external driving inputs to P1 and P2 used in Figure 4 were crafted using a multiscale noise generator that amalgamates slow and fast fluctuations. The noise was generated by combining two components: a slow drift and a fast fluctuation. The slow drift was produced using an autoregressive (AR(1)) process with coefficient αslow, yielding an effective timescale τslow = −Δt/ln(αslow), which captures long-term fluctuations. The fast component was generated as white noise, scaled by a standard deviation σfast, and then low-pass filtered with a 4th-order Butterworth filter having a cutoff frequency fcutoff, resulting in an effective timescale τfast = 1/(2πfcutoff). These two components were summed and offset by a base mean, and values were clipped to a minimum floor to prevent extremely small or negative outputs. Noise was generated over the simulation time vector with a sampling frequency fs = 1/Δt, and reproducibility was ensured by setting a fixed random seed.

Figure G1 shows an example of the multiscale noise generated and its power spectral density.

Generation of AM-Modulated Signals. To evaluate the impact of AM driving vs noise, we generated amplitude-modulated (AM) signals in the alpha band by applying nested modulation to a carrier tone.^120^ Formally, given a time array *t*, a carrier frequency *f_c_*, and two bandpass filters for modulators *m*_1_(*t*) and *m*_2_(*t*), we compute

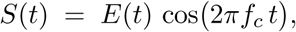

where the envelope E(t) is defined as

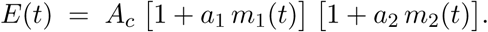

where a1 and a2 are the modulation indexes. In practice, m1(t) and m2(t) are obtained by bandpass filtering white noise within the specified frequency bands, and the filtered signals are then rescaled to the range [−1, 1]. The final AM-modulated signal S(t) is shifted to have zero mean. This nested structure allows the fast modulator to impose rapid changes in amplitude on the carrier, while the slow modulator introduces additional variations over longer timescales. In our simulation of priors, we set fc = 0, as we are interested in the modulated envelope only.

Driving the model with multiscale noise or AM signals produces similar CFC plots (healthy/nominal condition, see Figure G2 vs. Figure 4).

## H CFC and Power in MCI

In Figure H1, we provide SEC/EEC and related plots in the MCI condition (MCI) (CP V →P 2 = 300 and other parameters as described in the main text). The AD case (CP V →P 2 = 140) is not shown. Due to the drastic drop in gamma power, SEC and EEC are not meaningfully computed.

The CFC plots are consistent with the results presented in Figure 10. The positive correlation in SEC that facilitates error suppression via correct predictions is lost in MCI, where anticorrelation is present instead. This is coherent with the amplification of error in MCI through SEC in the case of a prediction matching the input (Figure 10e). Also, the lower EEC anticorrelation observed in MCI compared to healthy justifies the lower error suppression through EEC obtained in our simulations (Figure 10f).

### H.1 Effects of psychedelics on CFC for healthy, MCI, and AD conditions

In Figure H3, we provide CFC plots for the effects of psychedelics as described in the main text (increasing the synaptic gain of excitatory synapses targeting P1) in the nominal (“healthy”), MCI, and AD conditions. The AD case is not meaningful, as there is substantial power loss in the gamma and alpha bands. The MCI case suggests that psychedelics may help to improve cognitive function by recovering CFC, in particular SEC. Our previous work indicated that psychedelics can help oscillatory power in MCI,97 but here we can observe its positive impact in CFC, which impacts prediction error evaluation.

## I Bifurcation diagrams for MCI, AD, and Psychedelics

Here we prove the two-parameter bifurcation diagrams of the LaNMM model as a function of the external inputs φP1 and φP2 , as in Figure 2 of the main text, focusing on the effects of AD and psychedelics. The existence of limit cycles, i.e., oscillatory dynamics, is represented in colors: no limit cycle, steady-state (white), one limit cycle (light blue), and two limit cycles (dark blue). Within the region of two limit-cycles, the system can exhibit periodic motion, quasi-periodic, and chaotic dynamics.

We focus on how the parameter changes related to AD and psychedelics affect the oscillatory dynamics of the nominal (healthy) model, especially its ability to display multi-frequency activity. Figure I2 shows the effect of psychedelics, which is modeled by increasing the glutamatergic gain factor inputs to P1 (AP1 ). For the sake of comparison, in panel (a) we display the nominal case. We notice in panels (b)-(d) that as AP1 , the SNIC bifurcation (blue) gets displaced to the left, which means that oscillatory activity is initiated by smaller values of external inputs to P1. However, we also notice that the area where multiple limit cycles can coexist diminishes.

Figure I1 shows how impairing the connectivity between PV and P2 cells affects the dynamics of the healthy case, which is displayed in panel (a). Interestingly, decreasing the CP V →P2 to 300 yields a greater area of multi-frequency activity, which is due to the HB+ (grey) bifurcation being displaced to the left. By further decreasing CP V →P2 to the MCI case (c), we notice that the picture changes and HB+ gets displaced to the right, which causes a reduction in the area of multi-frequency activity. Finally, for the AD case (CP V →P2 = 140), the system only displays one limit-cycle, and only oscillates in the alpha band, in the region bounded by the SNIC and HB+ (purple) bifurcations as shown in panel (d).

Finally, the combined effects of psychedelics and MCI and AD are shown in Fig. I3. In panel (a), we show the two-parameter bifurcation diagram for the MCI case with psychedelics (CP V →P 2 = 300, AP1 = 4.5 mV). For these values, although the area of multi-frequency is smaller than in the nominal and MCI only case, we notice that the system is capable of exhibiting multifrequency activity for much smaller values of external inputs. The AD case with psychedelics (CP V →P 2 = 140, AP1 = 4.5 mV) is shown in panel (b), where we observe that the system does not exhibit oscillatory dynamics in the range of external inputs studied.

---

**Figure A1:**
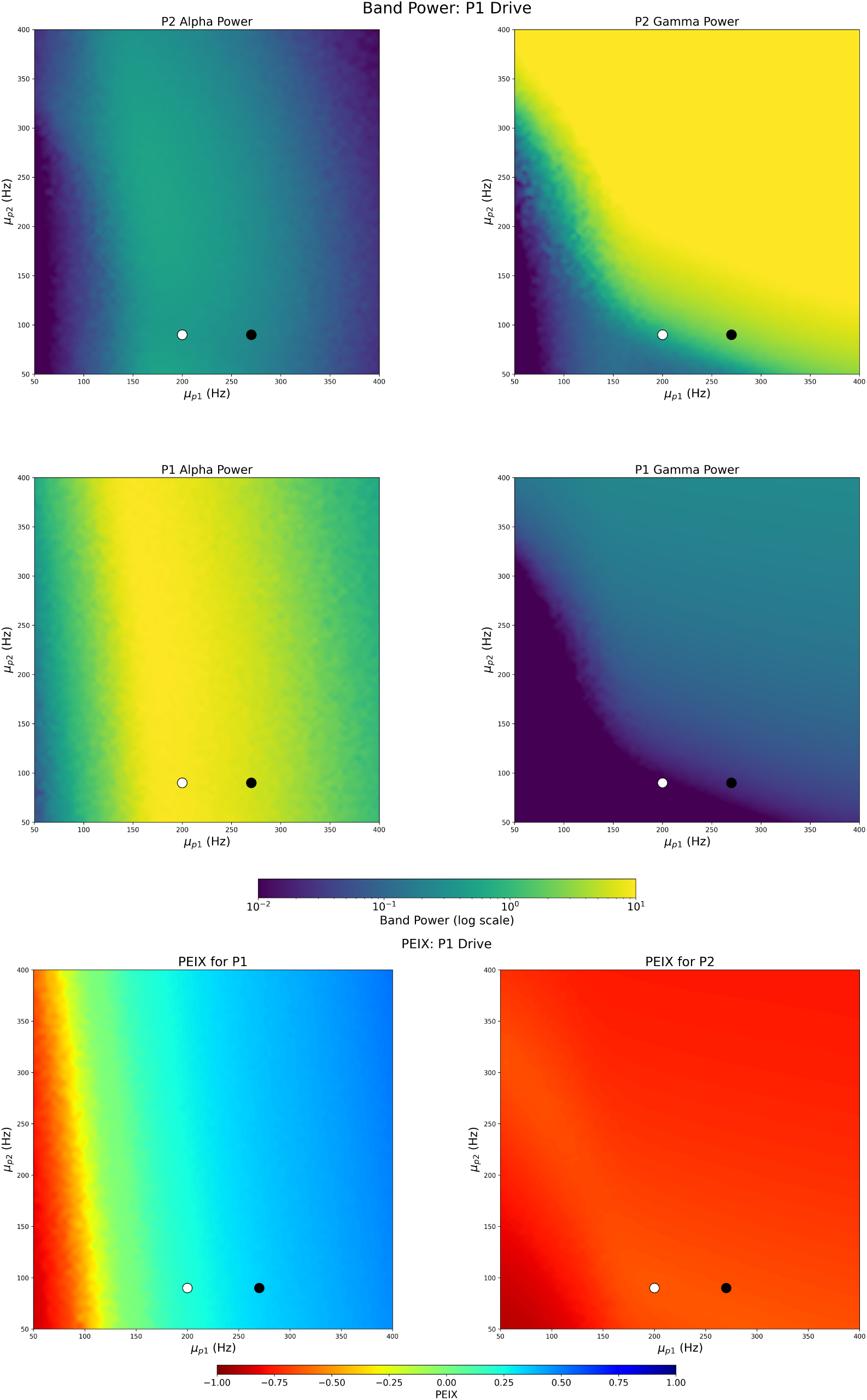
LaNMM band power for *P*_1_ and *P*_2_ populations for varying external mean drives *µ_P_*_1_ and *µ_P_*_2_ (stochastic drive on *P*_1_). The left column displays the alpha band power (8 − 12 Hz) for *P*_1_ and *P*_2_, while the right column shows the gamma band power (30 − 50 Hz) for each population. The white dot marks the mean input parameter values used in Sanchez-Todo el al. (2023),^105^ (*µ_P_*_1_ , *µ_P_*_2_ ) = (200, 90) Hz. The black dot indicates the values chosen for this study, (*µ_P_*_1_ , *µ_P_*_2_ ) = (270, 90) Hz. The bottom plot displays the PEIX sigmoid saturation indicator.

**Figure A2:**
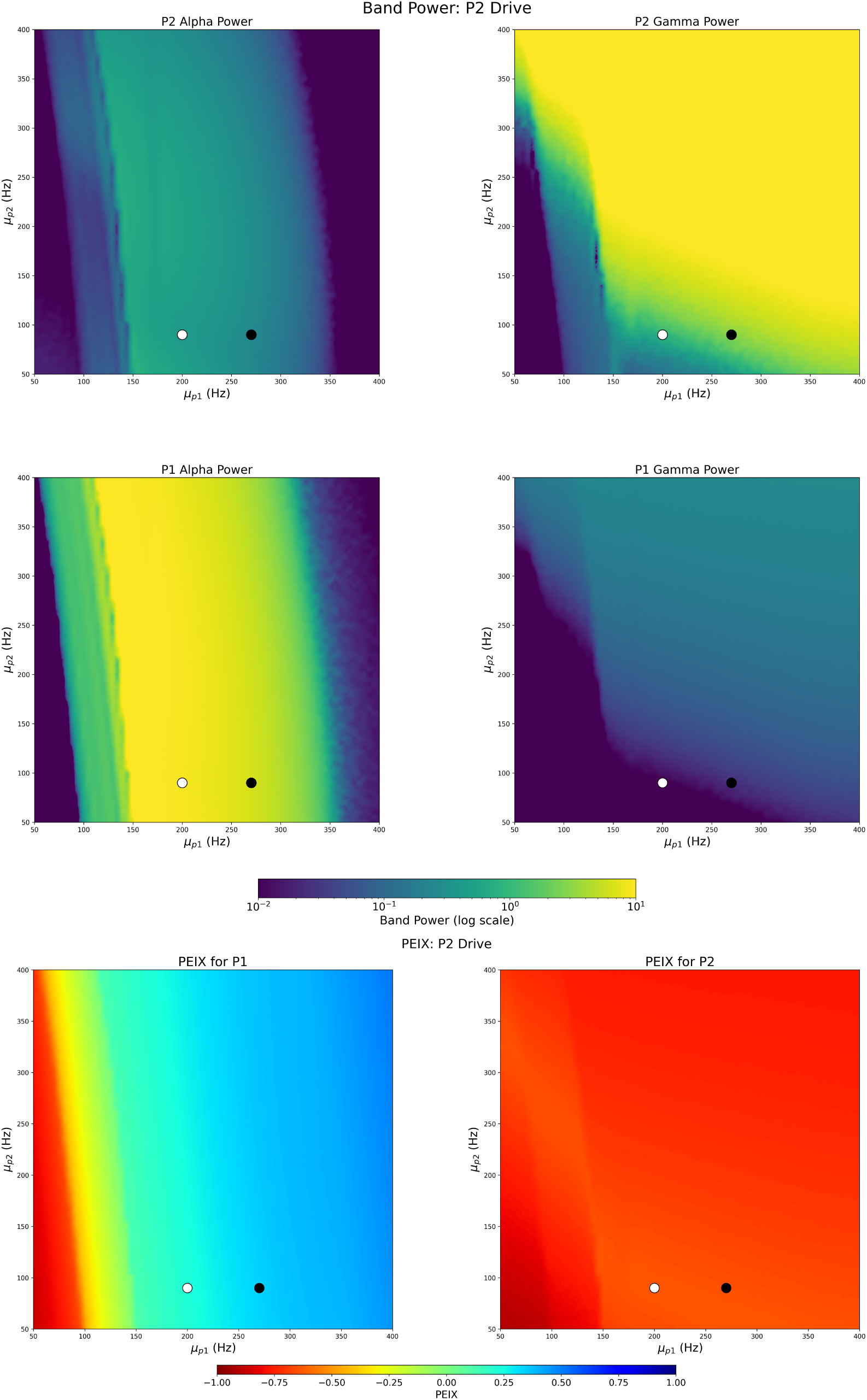
LaNMM band power for *P*_1_ and *P*_2_ populations for varying external mean drives *µ_P_*_1_ and *µ_P_*_2_ (stochastic drive on *P*_2_). The left column displays the alpha band power (8 − 12 Hz) for *P*_1_ and *P*_2_, while the right column shows the gamma band power (30 − 50 Hz) for each population. The white dot marks the mean input parameter values used in Sanchez-Todo et al. (2023),^105^ (*µ_P_*_1_ , *µ_P_*_2_ ) = (200, 90) Hz. The black dot indicates the values chosen for this study, (*µ_P_*_1_ , *µ_P_*_2_ ) = (270, 90) Hz, which is deeper into the quasi-periodic regime with stronger EEC (see Figures 4 and G2. The bottom plot displays the PEIX indicator. Appendix I provides bifurcation diagrams for the MCI, AD, and psychedelics cases.

**Figure B1:**
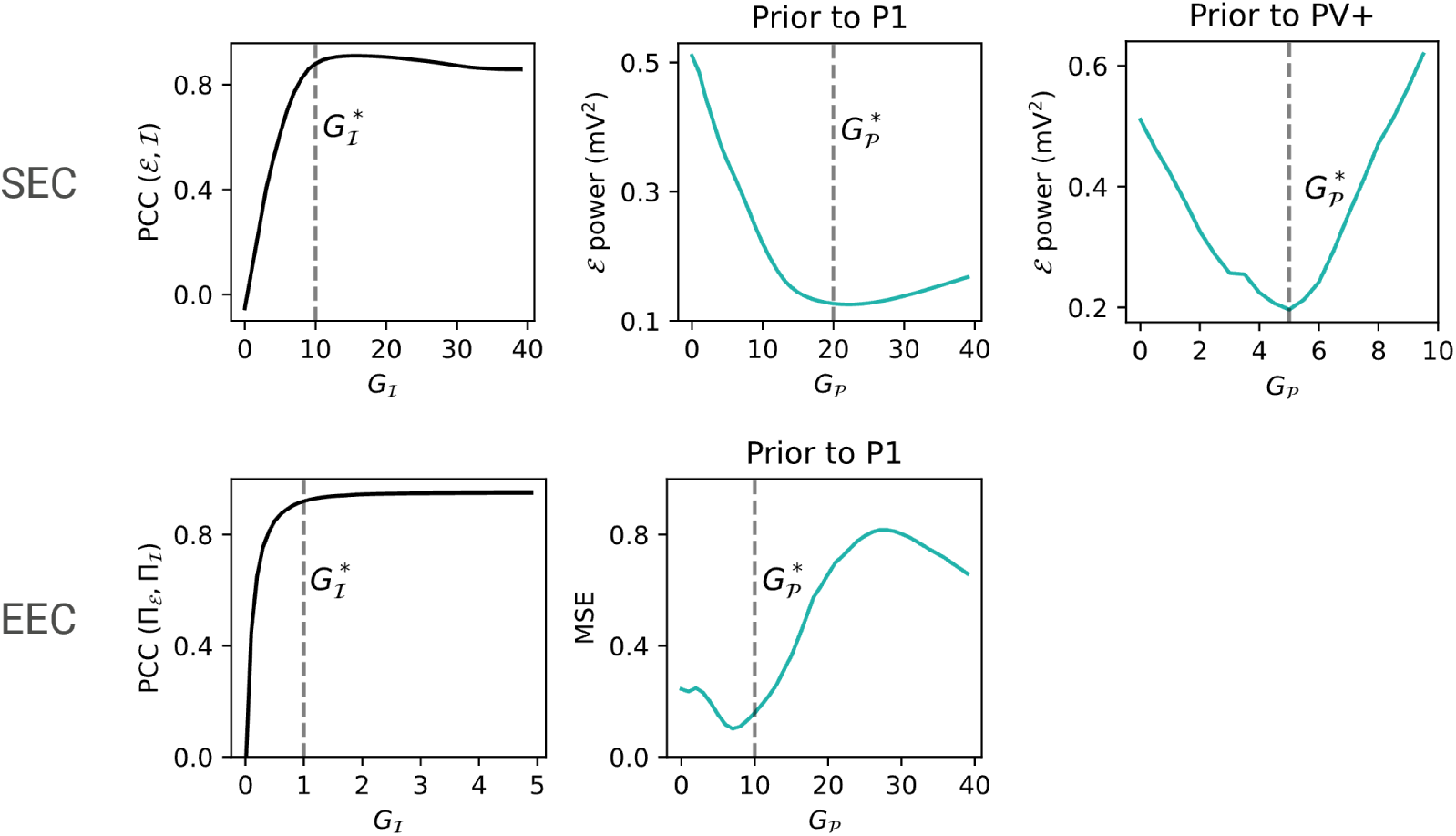
Selection of input and prior gains for SEC and EEC. Top and bottom panels show the gain optimization process for SEC and EEC, respectively. Left panels: Pearson correlation coefficient (PCC) between the LaNMM error signal E and the input message I as a function of input gain *G*_I_. The selected gain *G*^∗^ (dashed line) corresponds to the lowest value that ensures a high PCC. Middle and right panels: E power (top) and MSE (bottom) as a function of prior gain *G*_P_, showing a clear minimum when the prior matches the input. The selected prior gain ^∗^ (dashed line) approximates the value that minimizes error power.

**Figure C1:**
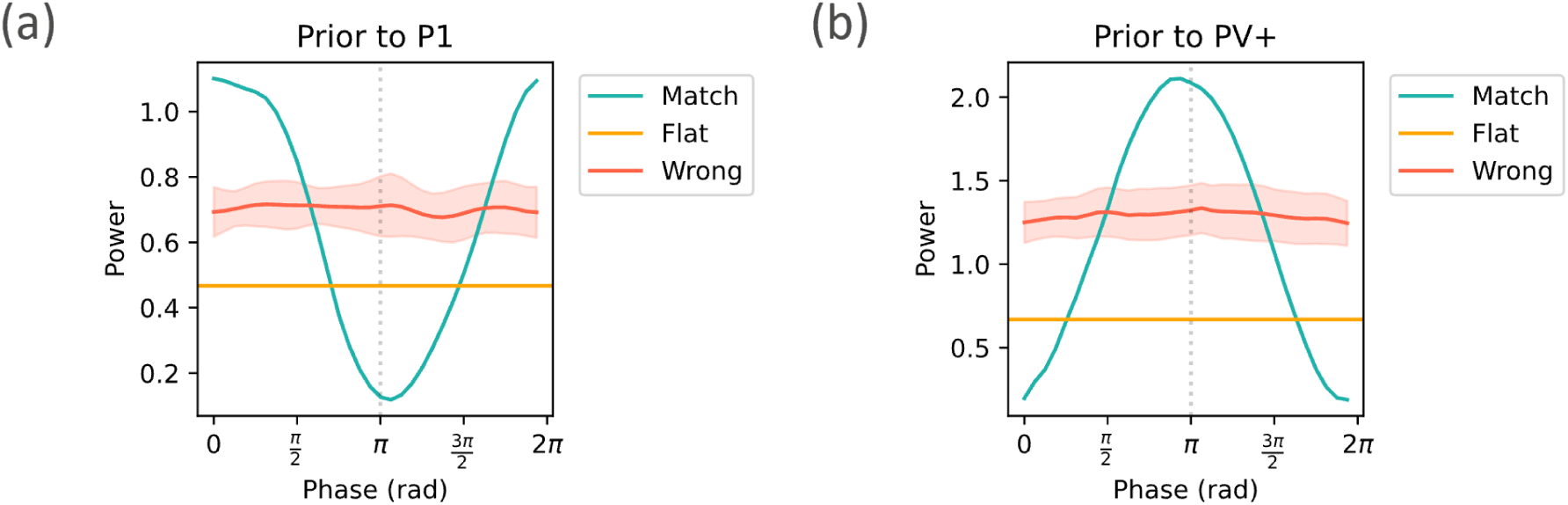
E power as a function of the phase shift of the prior with respect to the input signal. Different prior configurations are used: matching prior (“match”), wrong prior (“other”) and flat prior (“base”). (a) Prior to P1 condition. (b) Prior to PV condition.

**Figure E1:**
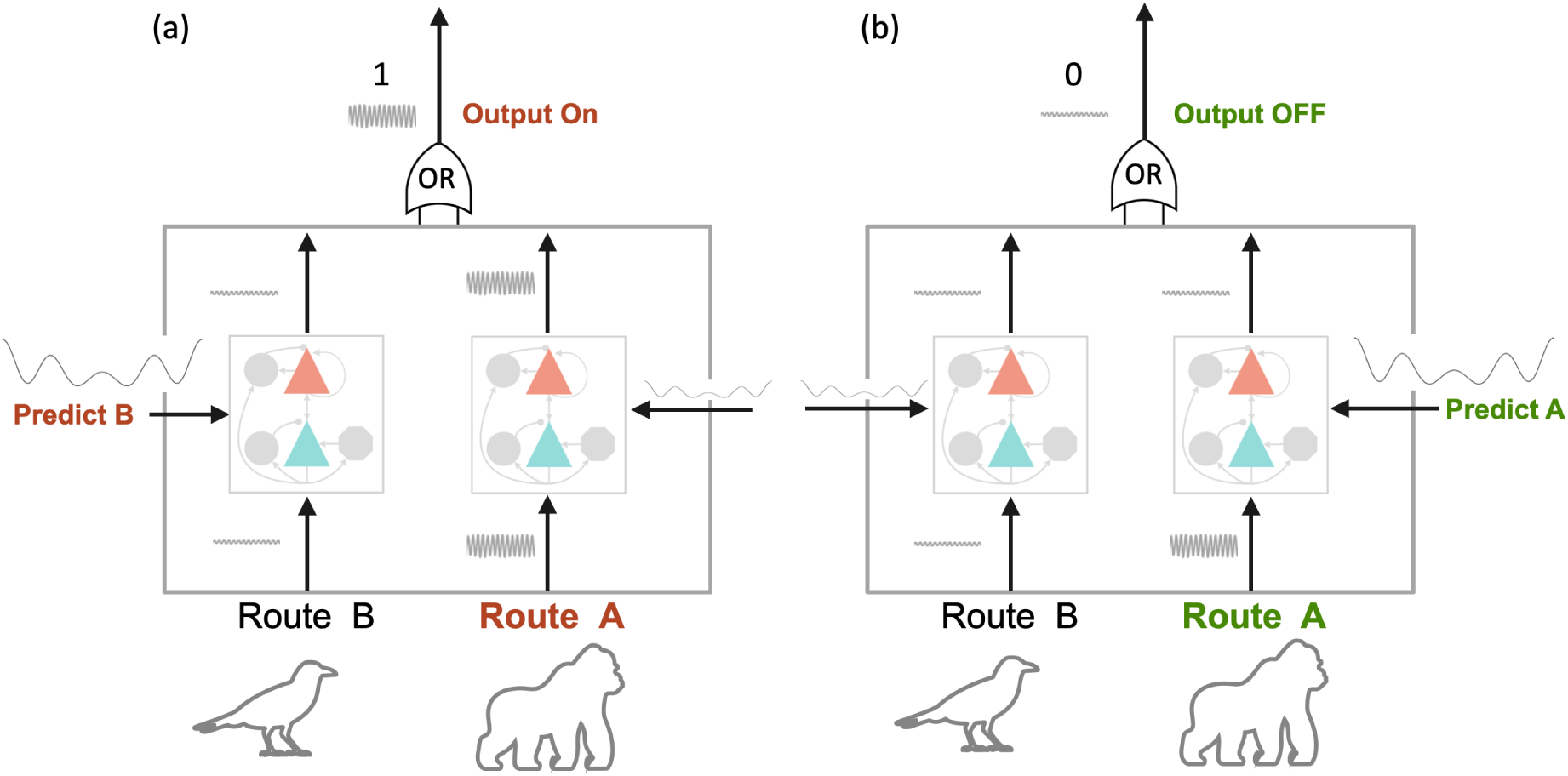
Predictive routing: prediction of the route of the stimulus: *information = route*. Bottom-up stimulus travels via route A or B (encoding 0 or 1, respectively), depending on the stimulus. In the depicted case, it travels via Route A (encoding a 1). The prediction aims to predict the route and inhibit information flow. Inhibition is implemented via EEC or SEC. (a) The route is incorrectly predicted (prediction on the left, encoding a 0), so the signal flows through (XOR(0,1)=1 on top, Output is ON). (b) The route is correctly predicted (Route 1 is predicted), and information flow is suppressed (XOR(0,0)=0).

**Figure F1:**
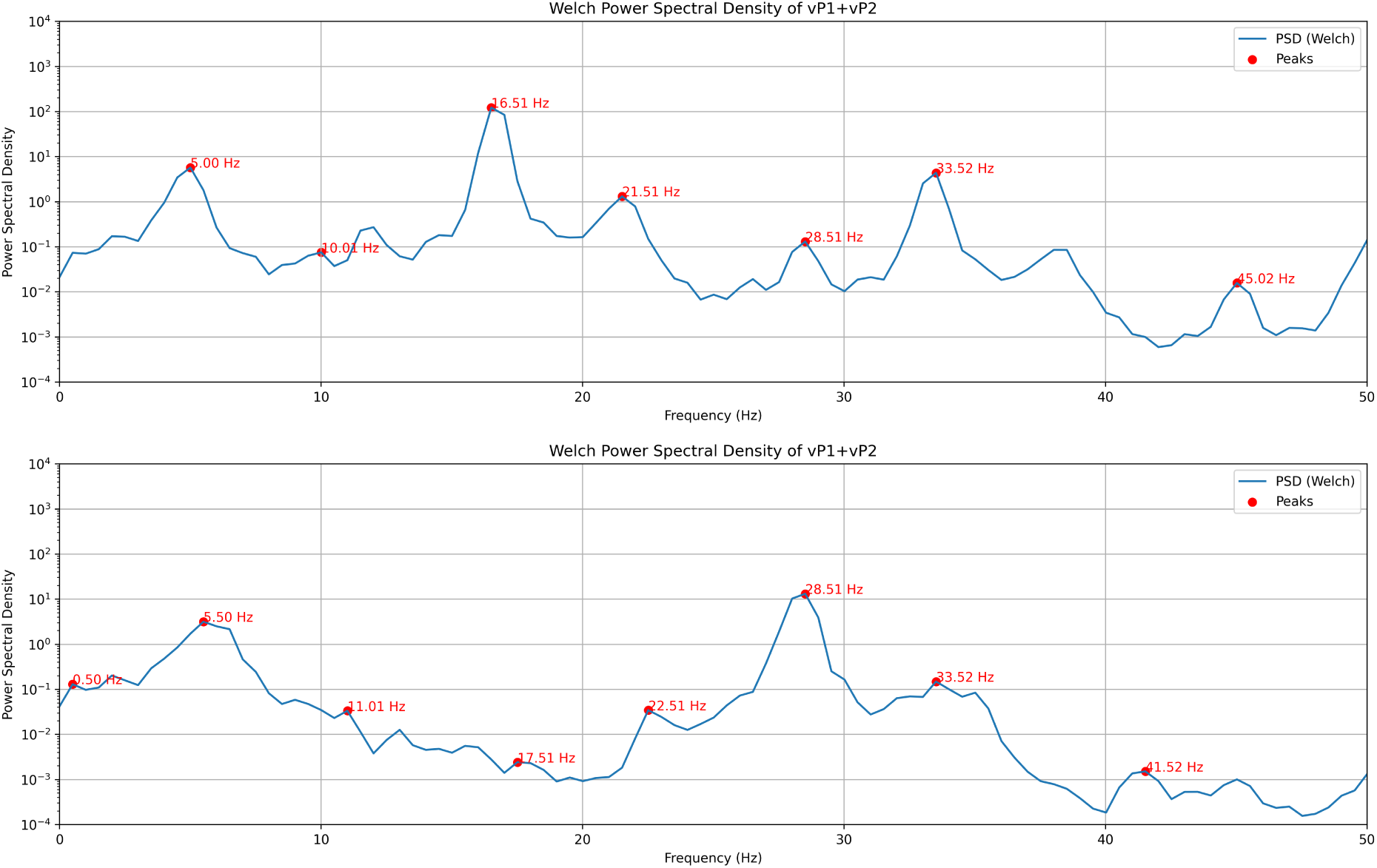
Examples from other LaNMM parameter configurations (only changing time constants).

**Figure G1:**
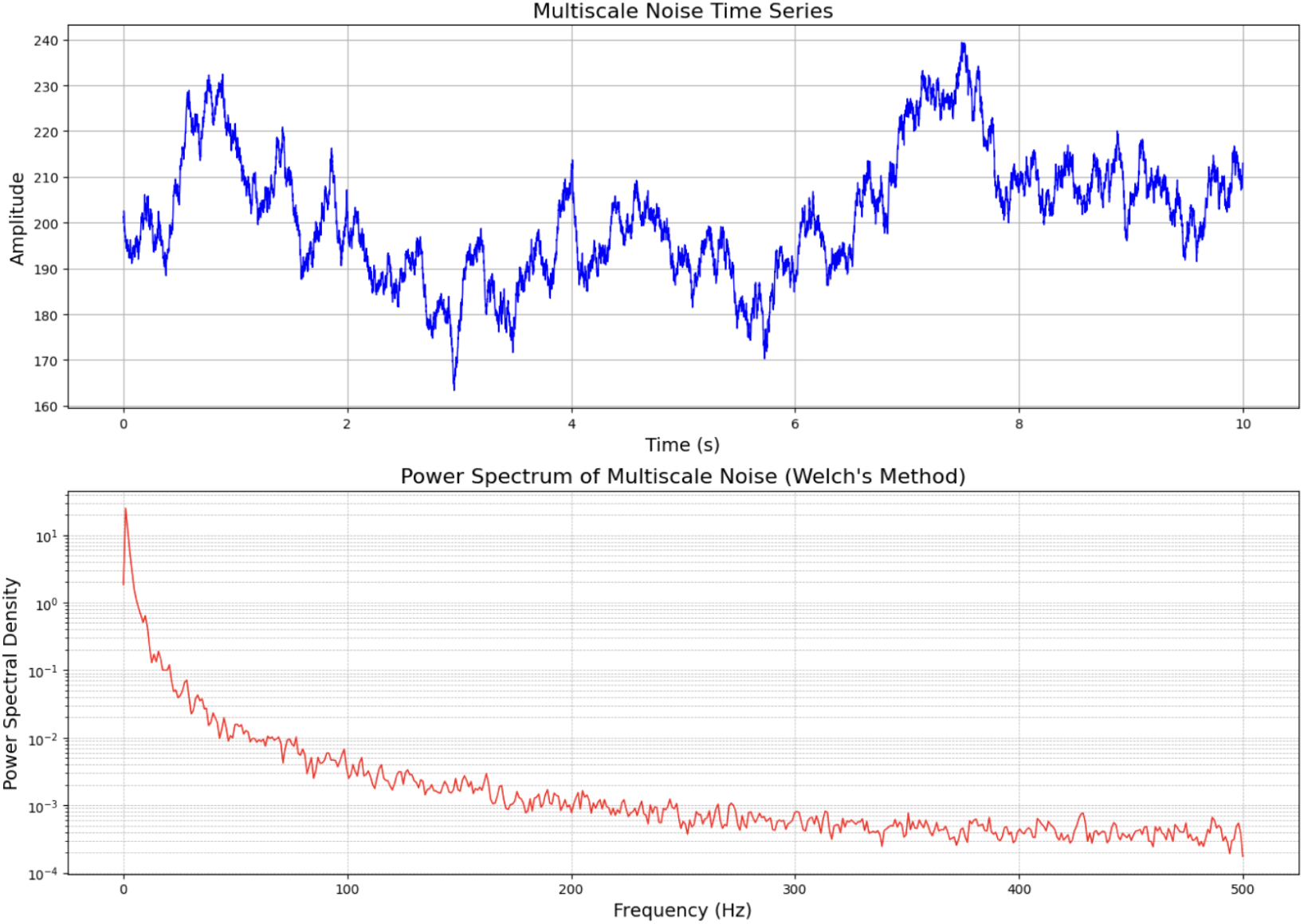
Example of multiscale noise (top panel) and its power spectral density (bottom panel).

**Figure G2:**
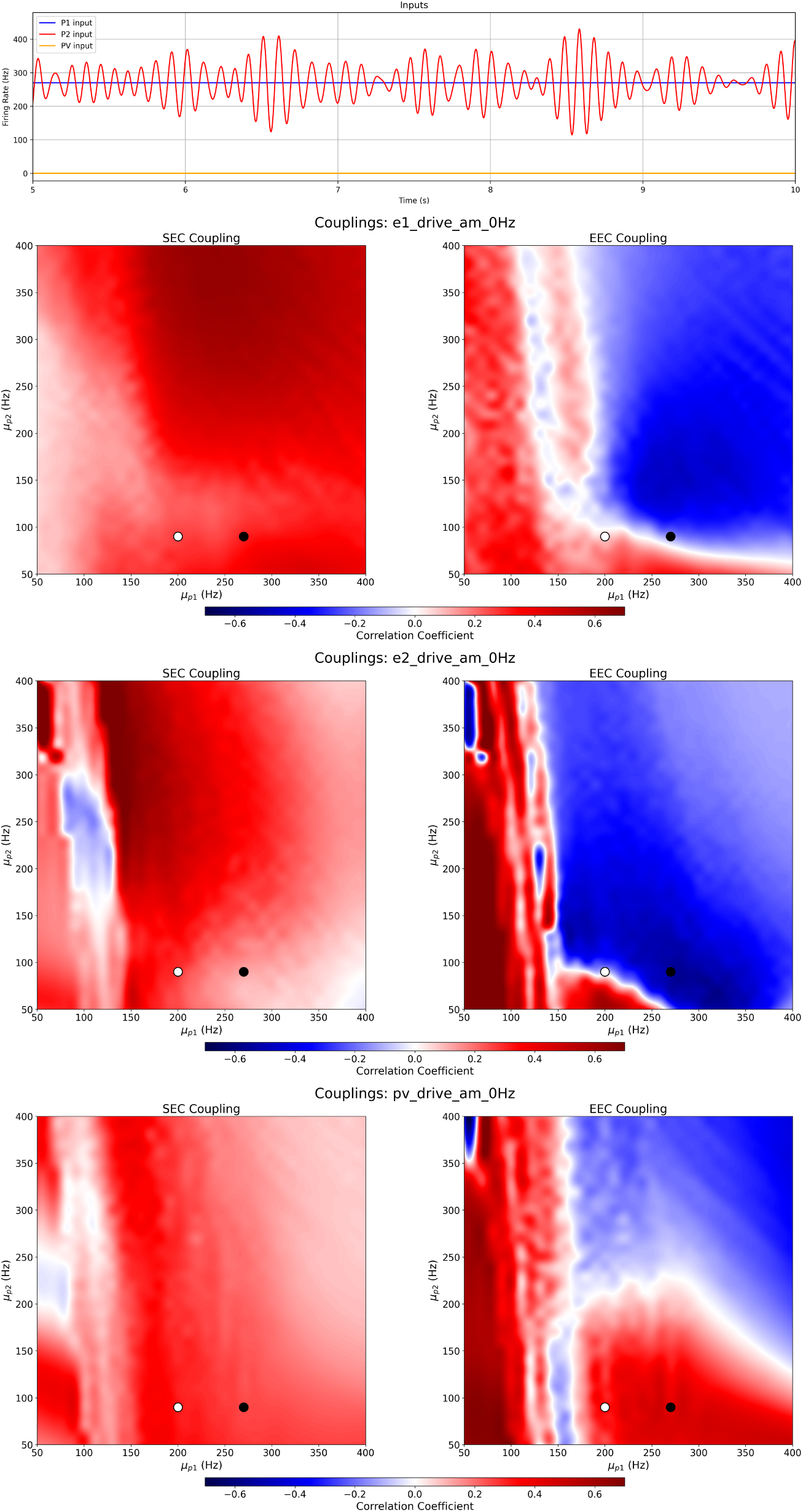
Top: sample modulation signal. Bottom: Couplings for P1, P2 and PV driving under AM modulation (0.5 Hz modulation of an 8-12 Hz band signal).

**Figure H1:**
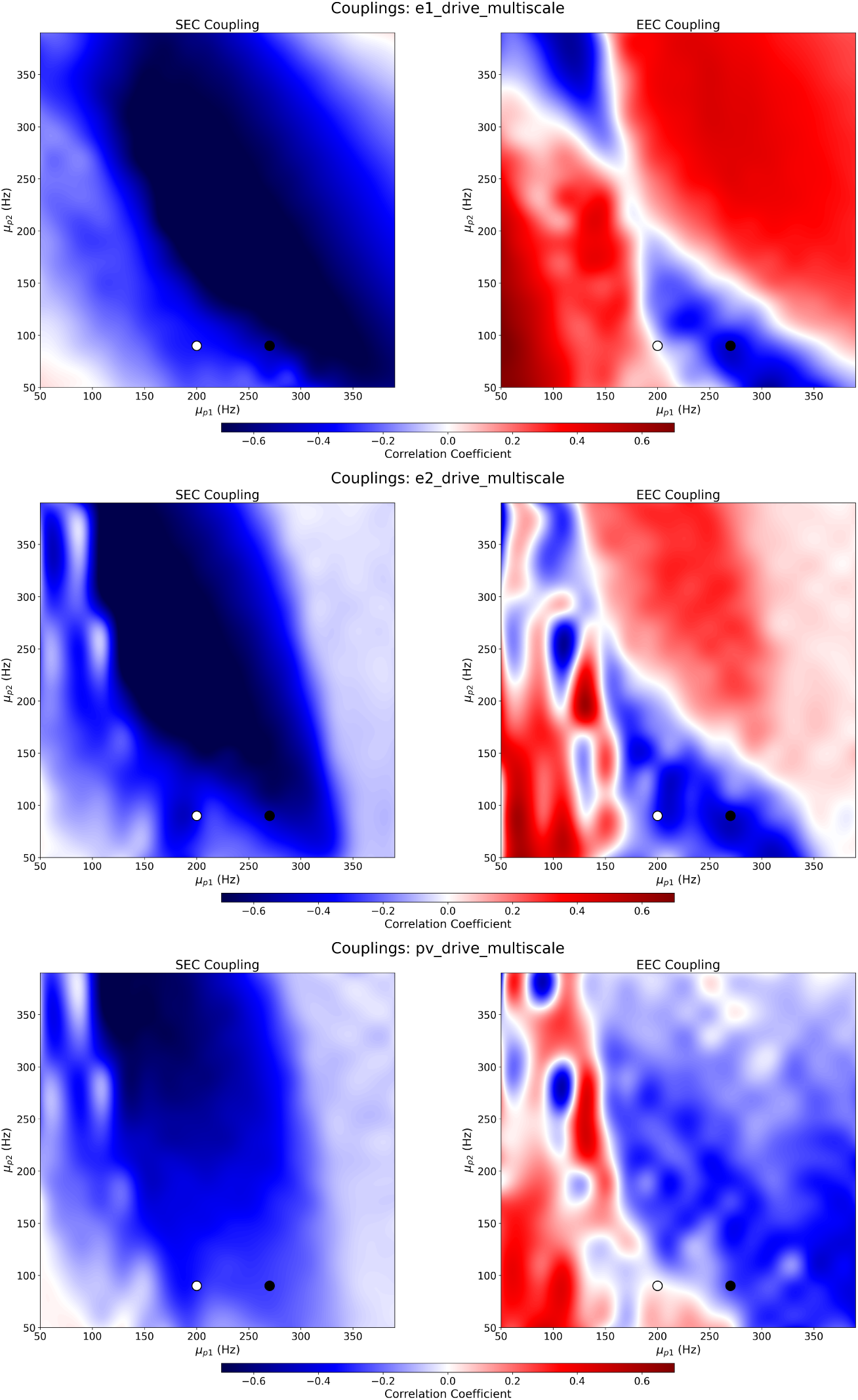
MCI condition: couplings under P1, P2 or PV multiscale noise drive. The negative SEC explains the increased prediction error in MCI — see Figure 10.

**Figure H2:**
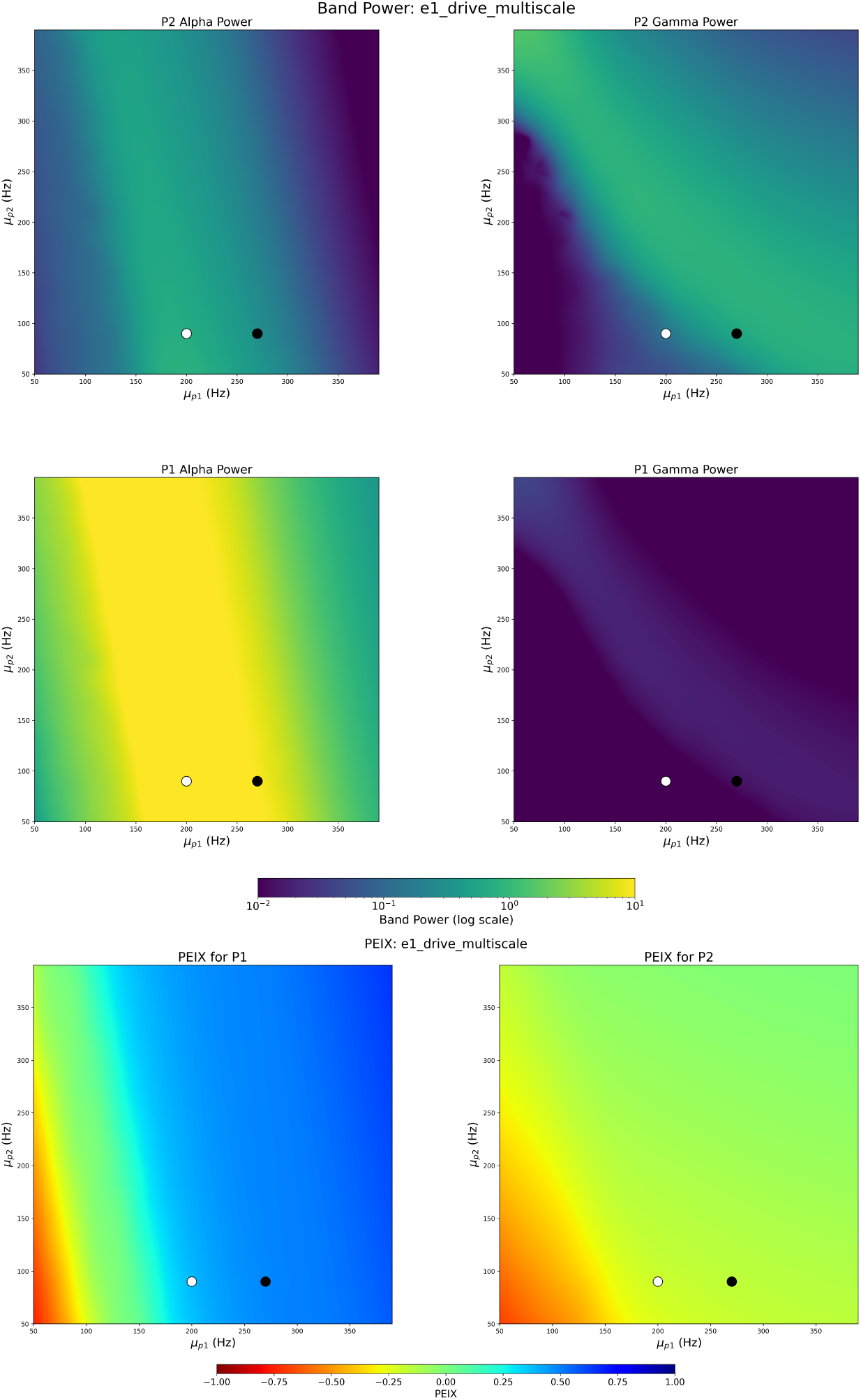
MCI condition: Power and PEIX heatmaps.

**Figure H3:**
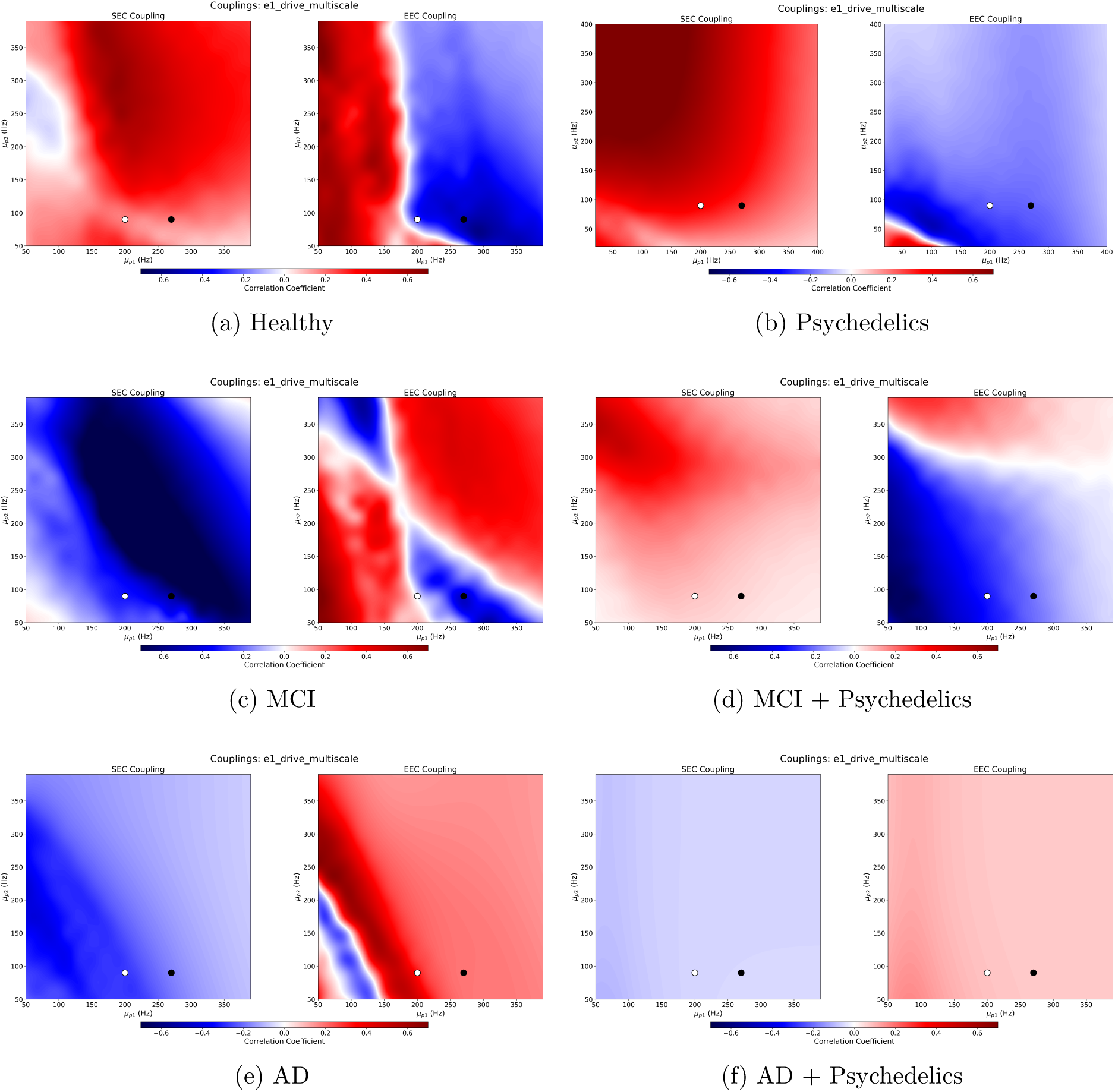
(a, b) CFCs in healthy vs psychedelics condition. (c, d) MCI vs MCI+Psychedelics, showing improvement toward healthy SEC. (e, f) AD vs AD+Psychedelics, not very meaningful as gamma activity is missing in the model.

**Figure I1:**
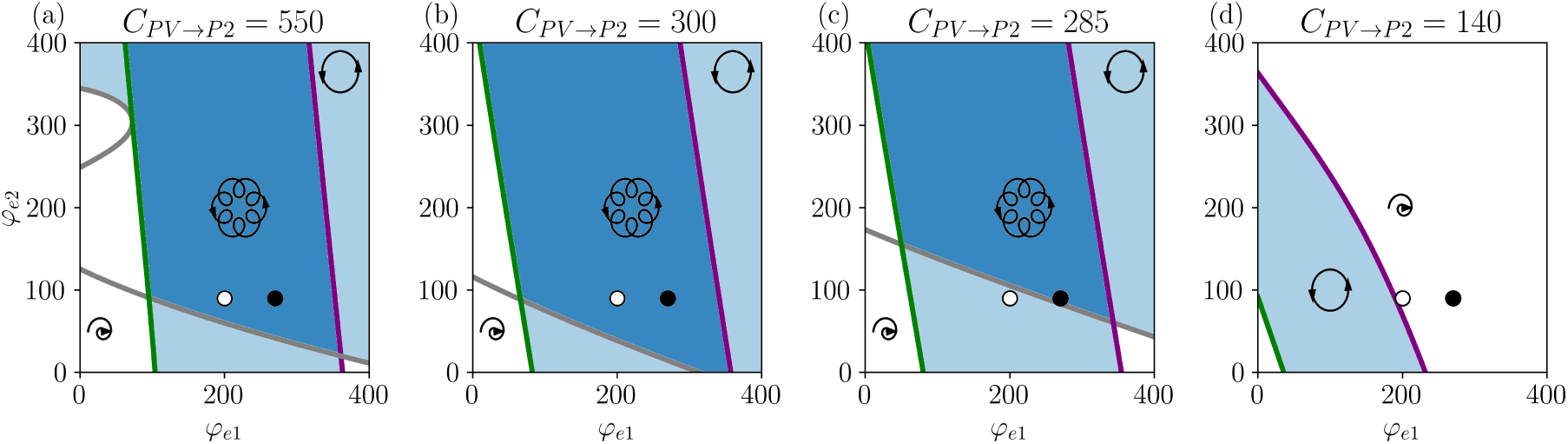
Bifurcation diagrams for AD case. From right to left, healthy PV connectivity to MCI to AD. The multifrequency area (dark blue) gets progressively smaller until it disappears.

**Figure I2:**
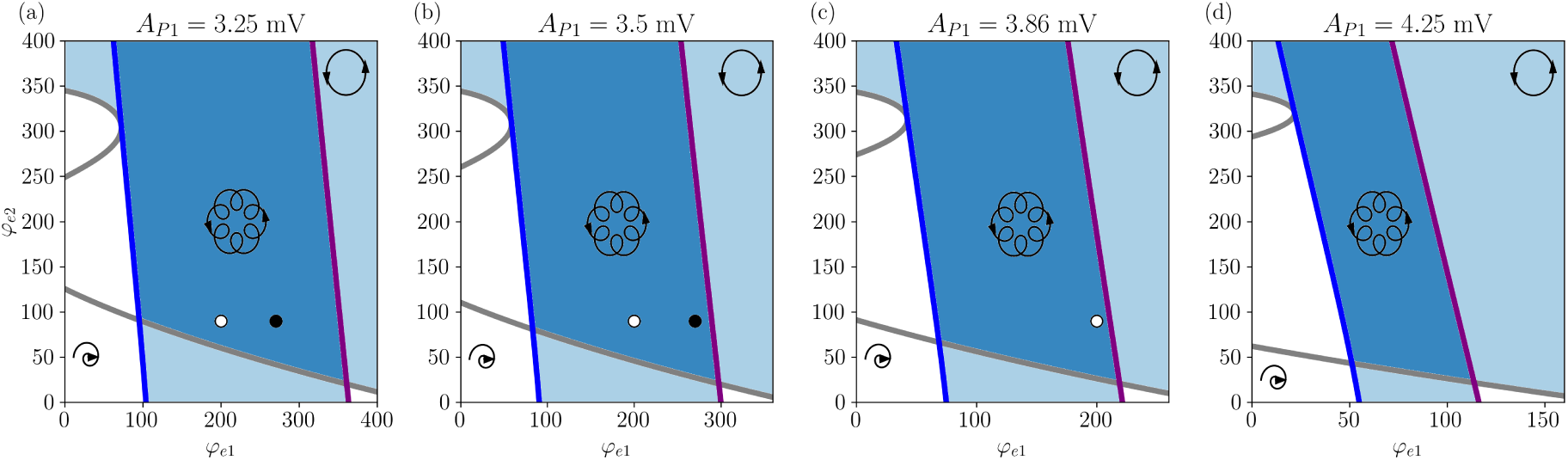
Bifurcation diagrams for the Psychedelics case on the nominal (healthy) model. The factor *A*_P1_ denotes the glutamaterigic synaptic gain of connections to P1.

**Figure I3:**
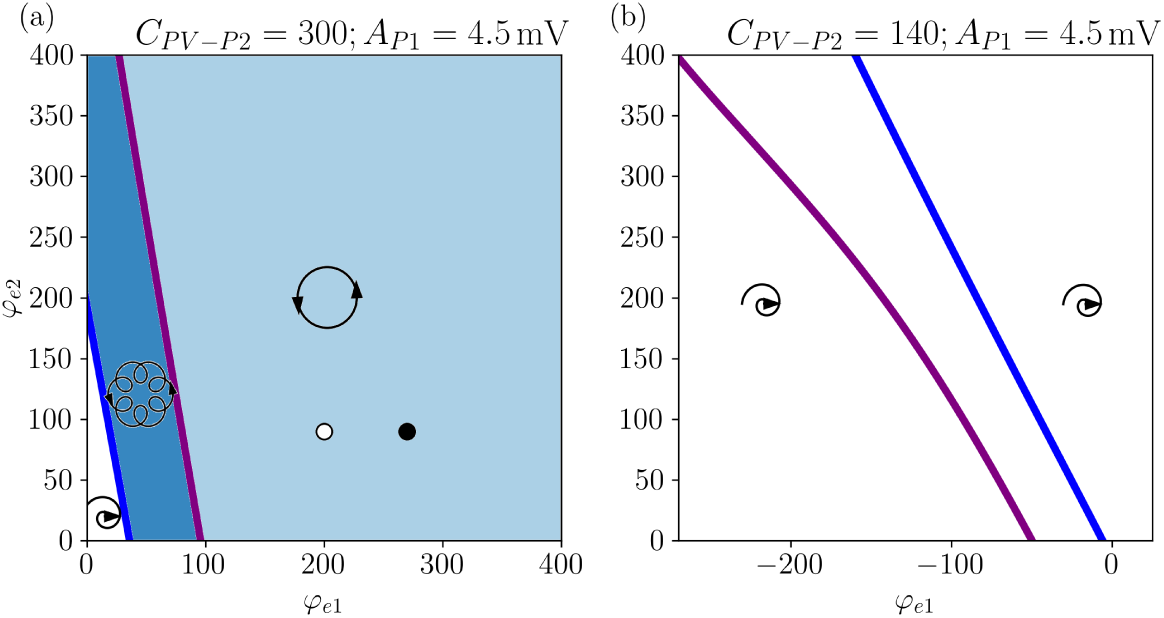
Bifurcation diagrams of AD and MCI under psychedelics.

## Notes

### Competing Interest Statement

GR, ELS, FC, and RST work for Neuroelectrics. GR is a co-founder.

### Summary of Updates

Substantial modifications to streamline the text, plus new appendix sections / bifurcation diagrams.

